# Histone Deacetylase Inhibitor Largazole Deactivates A Subset of Superenchancers and Causes Mitotic Chromosome Mis-alignment by Suppressing SP1 and BRD4

**DOI:** 10.1101/2025.01.29.635612

**Authors:** Gilson J. Sanchez, Zeyu Liu, Samuel Hunter, Quanbin Xu, Jessica T. V. Westfall, Graycen E. Wheeler, Cathryn Toomey, Dylan Taatjes, Mary Allen, Robin D. Dowell, Xuedong Liu

**Author notes:** Correspondence and Lead Contact (X.L.). These authors contributed equally (Gilson J. Sanchez, Zeyu Liu).

## Abstract

Histone deacetylase inhibitors have been investigated as potential therapeutic agents for cancer and other diseases. HDIs are known to promote histone acetylation, resulting in an open chromatin conformation and generally increased gene expression. In previous work, we reported that a subset of genes, particularly those regulated by superenhancers, can be suppressed by the HDAC inhibitor largazole. To elucidate the molecular mechanisms underlying gene repression by largazole, we conducted transposase-accessible chromatin sequencing, ChIP-seq, and RNA-seq studies. Our findings revealed that while largazole treatment generally enhances chromatin accessibility, it selectively decreases the accessibility of a subset of superenhancer regions. These genomic regions, showing the most significant changes in the presence of largazole, were enriched with transcription factor binding motifs for SP1, BRD4, CTCF, and YY1. ChIP-seq analysis confirmed reduced binding of BRD4 and SP1 at their respective sites on chromatin, particularly at superenhancers regulating genes such as ID1, c-Myc and MCMs. Largazole exerts its effects by inhibiting DNA replication, RNA processing, and cell cycle progression, partially through the suppression of SP1 expression. Depletion of SP1 by shRNA mimics several key biological effects of largazole and increases cellular sensitivity to the drug. Specific to cell cycle regulation, we demonstrated that largazole disrupts G/M transition by interfering with chromosome alignment during metaphase, a phenotype also observed with SP1 depletion. Our results suggest that largazole exerts its growth-inhibitory effect by suppressing BRD4 and SP1 at super-enhancers, leading to cytostatic responses and mitotic dysfunction.

## INTRODUCTION

Acetylation of histones alters the compaction of chromatin, making it more accessible to transcription factors and the transcriptional machinery. The relaxed chromatin conformation allows regulatory proteins to bind to specific DNA sequences more easily, facilitating gene activation. The balance between histone acetyltransferases (HATs) and histone deacetylases (HDACs), which add or remove acetyl groups respectively, is crucial for maintaining proper histone acetylation levels and cell-type or tissue specific gene expression. Deregulation of histone acetylation due to the hyper-or aberrant activity of HDACs has been observed in human tumors. Because of their roles in tumorigenesis, HDACs have been identified as potential targets for cancer therapies ^1^. As a result, four HDAC inhibitors (Vorinostat, Belinostat, Romidepsin and Panobinostat) have been approved by FDA for treating hematological cancers ^2^. Even though histone deacetylase inhibitors (HDIs) are used clinically, the molecular mechanisms by which these inhibitors suppress tumor growth beyond histone acetylation remain debatable. Some of mechanisms that have been proposed include: 1) activation of tumor suppressor genes and the repression of oncogenes such as Myc, NF-kB, and STAT3, leading to a decrease in cell proliferation and survival; 2) induction of cell cycle arrest by downregulating cyclins thereby inhibiting cyclin-dependent kinase (CDKs); 3) induction of apoptosis by upregulating pro-apoptotic genes, such as Bax and Bak, while downregulating anti-apoptotic genes, such as Bcl-2; 4) inducing DNA damages by suppression of DNA repair pathways; 5) Immune modulation by elevating the expression of major histocompatibility complex (MHC) and immunogenicity of tumors. A unifying theme of these proposed mechanisms is that HDIs work by up or down regulating a subset of genes that are relevant to tumorigenesis process ^1,2^. However, the precise mechanisms of HDI-induced transcriptional activation or suppression are still yet defined.

Even though HDAC inhibition causes global increase in histone acetylation and chromatin accessibility, the outcomes of gene transcription can be either up or down regulated. Acetylation of a subset of promoters can be decreased with the treatment of HDIs leading transcriptional suppression. We and others reported that increased acetylation in the gene body affects transcriptional elongation resulting in RNA polymerase pausing ^3–5^. Also, we reported that superenhancers (SEs) are preferentially targeted by HDIs, which correlate with decreased expression of SE-regulated genes such as c-Myc ^5^. One explanation for SE perturbation by HDIs is that they causes genomic redistribution of SE activator BRD4, which specifically recognizes and binds to acetylated lysine residues on histones, particularly H3K27ac, from SE regions to other areas of polyacetylated chromatin (Slaughter et al. 2021). However, not all SE are deactivated by HDIs. The mechanisms why some SEs are more sensitive to HDI inhibition than others remain unclear.

Largazole, a natural product that was originally isolated from the marine cyanobacterium, belongs to the class of cyclic depsipeptides and exhibits potent inhibitory activity against Class I and Class IIb HDAC enzymes ^6,7^. Largazole treatment induces cell cycle arrest and apoptosis in various cancer cell lines *in vitro* and tumor suppression *in vivo* ^8^. Largazole derivative, OKI-179, is being evaluated for anticancer activity in phase II clinical trials ^9–11^. Our previous results showed that largazole causes remodeling of numerous enhancer elements by modulating H3K27ac and suppression of a subset of SEs in a dose-dependent manner ^5^. Here we investigate the effects of largazole treatment on chromatin accessibility and transcription factor binding to SE elements at the genome level globally. We find that HDAC inhibition by largazole results in robust increase in chromatin accessibility and loss of BRD4 binding in selective SE elements. Transcription factor binding site enrichment analysis for genomic regions whose accessibility is regulated by largazole treatment revealed that transcription factor SP1 binding sites are significantly under-represented. ChIP-seq analysis shows that loss of SP1 binding in promoters and enhancers, particularly, SEs where BRD4 binds. Furthermore, we uncovered that a new mechanism for largazole-induced cell cycle arrest by perturbing chromosome alignments through suppression of SP1.

## RESULTS

### HDAC inhibition by largazole results in displacement of BRD4 from enhancer and transcription initiation sites

We previously established that treatment of HCT116 cells with the HDAC inhibitor largazole leads to the de-acetylation of Lys–9 and Lys–27 on histone H3 at a subset of regulatory regions leading to the inactivation of hundreds of enhancer elements ^5^. BRD4 is known to occupy proximal gene promoters and enhancers and mediate transcriptional activation ^12^. We therefore investigate whether HDAC inhibition may disrupt BRD4 binding to promoters and enhancers by BRD4 ChIP-seq in HCT116 cells. JQ1, a well-established inhibitor of BRD4 ^13^, was used as a control. Cells treated with (i) vehicle, (ii) largazole, (iii) JQ1, (iv) JQ1+largazole for 16 h were processed for BRD4 ChIP-seq studies. Shown in Figure 1A is a representative snapshot of BRD4 peaks at NEAT1 and MALAT1 promoters under the four experimental conditions in comparison with input genomic DNA. In total, 17,216 BRD4 binding sites identified under basal conditions were found to be at transcription initiation sites in gene promoters (37.2%) and enhancer elements (24.1%) (Figure 1B). In agreement with previous reports ^13^, treatment of HCT116 cells with JQ1 abolished most BRD4 interactions with chromatin, and only 6,233 peaks persisted after treatment (71.2% reduction). Notably, largazole treatment alone is capable of disrupting the association of BRD4 with chromatin resulting in 8,196 peaks (54.7% reduction) and similarly to the effect of JQ1, but to a lesser extent. Largazole primarily disrupts BRD4 peaks localized within enhancers and proximal gene promoters (Figure 1B). With combined treatment with JQ1 and largazole resulted in the mapping of 7,016 BRD4 peaks (59.1% reduction) with a more drastic reduction of BRD4, as 497 gene promoters displayed complete eradication of BRD4 signal under the combined treatment relative to either single treatment as seen in NEAT1 and MALAT1 promoters (Figure 1A and 1B).

**Figure 1.**
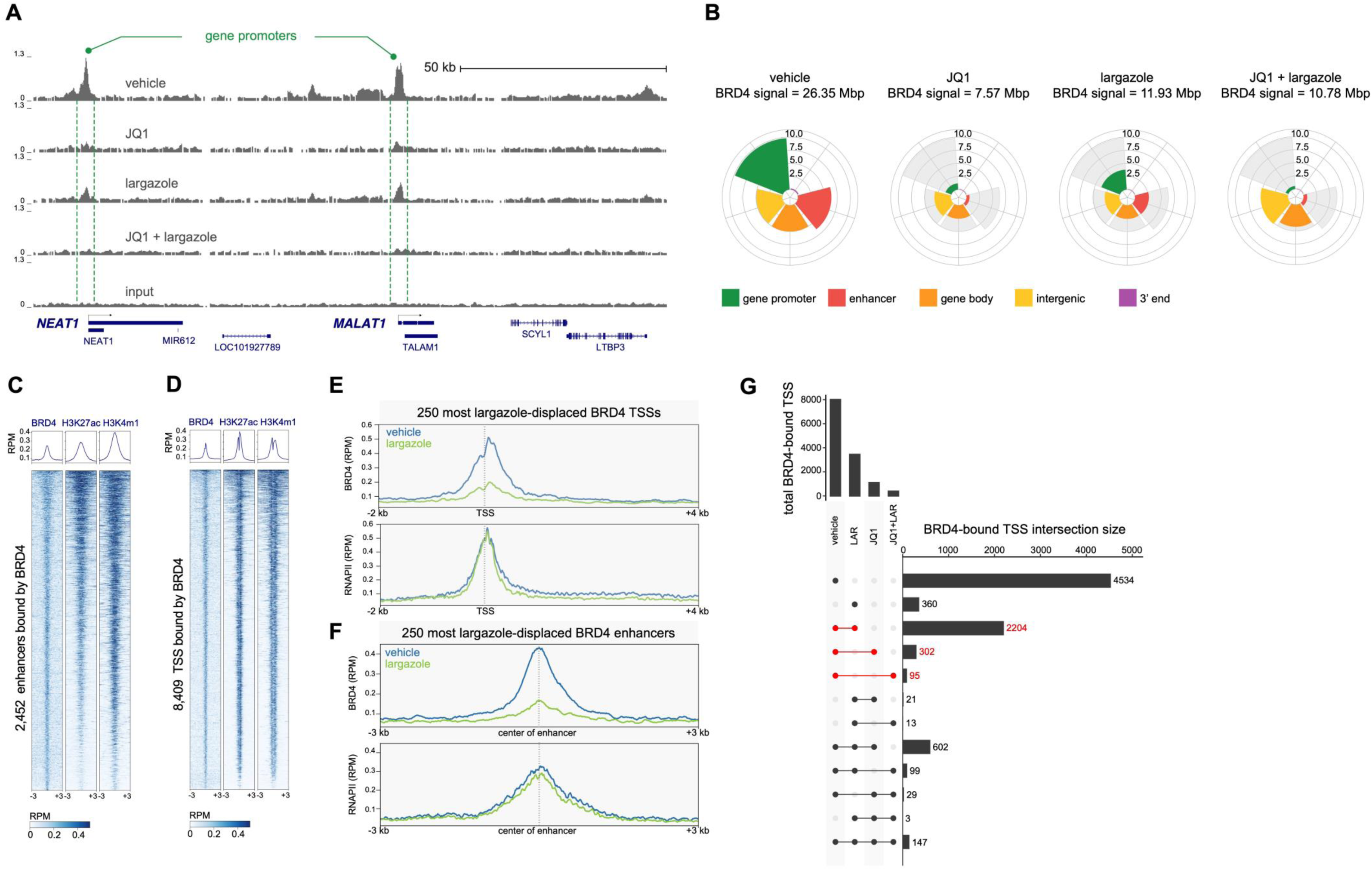
Largazole, JQ1 or their combination displace BRD4 from a subset of enhancers and proximal promoters. (**A**) Genome browser visualization of the NEAT1 and MALAT1 gene loci depicting BRD4 ChIP-seq signal derived from HCT116 cells under four treatment conditions: 10 µM JQ1, 75 nM largazole, the combination of JQ1 and largazole, or the vehicle control (DMSO). (**B**) Polar charts showing the total distribution (in Mbp) of BRD4 ChIP-seq signal (determined by MACS2) across four treatment conditions. BRD4 distribution is represented as bar plots, with axes labeled in megabase pairs (Mbp) and color-coded for five genomic categories: gene promoters (green), enhancer regions (reds), gene body (orange), intergenic locations (yellow), and 3′ ends (purple). The grey-shaded regions denote BRD4 signal under control conditions. (**C**) and (**D**) ChIP-seq read counts (Reads Per Million mapped, RPM) for BRD4-bound genomic elements, centered on each MACS2-defined BRD4 peak summit and extending ±3 kb. Panel **C** shows 2,452 enhancers, and panel **D** shows 8,409 transcription start sites (TSSs). (**E**) and (**F**) Change in BRD4 (top) and RNA Pol II (bottom) accumulation within genomic regions following largazole treatment. BRD4 reads are normalized by both the length of the genomic element (in kilobases) and the total number of mapped reads (RPKM). Genome-wide ChIP-seq profiles of BRD4 (top) and RNA Pol II (bottom) for the 250 most largazole-displaced TSS (**E**) or enhancers (**F**) centered ±3 kb around the BRD4 peak summits or RNA Pol II summits. Signals are shown for vehicle (blue) and largazole (green). (**I**) UpSet plot summarizing the intersecting distribution of BRD4 across 8,409 TSS regions in response to four distinct cellular conditions.

To further illustrate the effect of largazole on BRD4 occupancy at enhancers and transcription start sites (TSS), we plotted the peak intensity ratios of 2,452 BRD4-bound enhancers marked with H3K27ac and H3K4me1 in largazole-treated cells relative to vehicle-treated controls (Figure 1C and 1D). Among these, the top 250 largazole-displaced enhancers showed a clear reduction in BRD4 signal, accompanied by a slight decrease in Pol II signals (Figure 1F). In contrast, at the top 250 largazole-displaced TSS regions, Pol II signals in the gene body were notably lower compared to the signals observed at the TSS (Figure 1E). This finding aligns with previous reports that HDAC inhibitor (HDI) treatment can induce Pol II pausing in a subset of genes, thereby affecting transcription elongation ^3,5^. These results suggest that largazole treatment significantly alters BRD4 occupancy at enhancers and TSSs, with distinct effects on Pol II distribution and transcription dynamics.

To determine how different treatments affect BRD4 association at TSSs and promoters, we utilized an UpSet plot to visualize the overlap of BRD4 binding sites under various conditions. Of the 8,012 BRD4 binding sites identified at TSSs across all conditions, 4,534 are exclusive to the vehicle condition, indicating that 3,478 BRD4 sites are altered by single or combined treatments. In contrast, only 360 sites are exclusive to the largazole-only treatment. The largest overlap is between vehicle and largazole, comprising 2,204 BRD4 sites, whereas only 302 sites overlap between vehicle and JQ1, and just 95 sites overlap between vehicle and JQ1 plus largazole (Figure 1G, connected red dots). These findings suggest that, in terms of BRD4 loss at TSSs, largazole and JQ1 do not fully overlap. Moreover, the combined JQ1 plus largazole treatment leads to a more pronounced eviction of BRD4 from its cognate sites. A similar pattern is observed at enhancers (Supplemental Figure 1A). Overall, our results reveal that largazole alone can displace BRD4 from transcription initiation sites, while the combined treatment maximizes BRD4 displacement from proximal gene promoters, implying both overlapping and non-overlapping mechanisms for BRD4 recruitment to chromatin

### HDAC inhibition by largazole disrupts SEs and expression of SE-associated genes

BRD4 is a well-characterized regulator of SEs which are responsible for the expression of hundreds of transcripts including many proto-oncogenes ^13,14^. Since we previously demonstrated that largazole disrupts SEs and SE-dependent activity, we therefore investigated the impact of largazole or JQ1 or combined treatment on SEs. To determine drug-dependent changes on BRD4 occupancy along SEs, we first used the SE set (n = 385) generated based on H3K27ac signal from HCT116 cells by the dbSUPER database ^15^. The density of BRD4 occupancy was established by tabulating the number of ChIP-seq reads within the boundaries of individual SE and normalized to the length of the SE region and to the total number of mapped fragments from the corresponding library (RPKM). We found BRD4 accumulation at SE is affected differently by single JQ1 or largazole drug treatments (Figure 2A). Consistent with the meta-analysis of the BRD4 ChIP-seq data, JQ1 is more efficient at displacing BRD4 from SE locations relative to the effect of largazole. For example, the SE located ∼400 kb upstream of the Myc locus is highly occupied by BRD4 (1.65 RPKMs) but highly depleted after JQ1 treatment (0.50 RPKMs). Exposure to largazole reduces the accumulation intensity (1.05 RPKMs) but fails to fully evict BRD4 (Figure 2A and 2B). Interestingly, the combined treatment results in maximal displacement of BRD4 from the vast majority of SEs where the mean ChIP-seq signal is significantly lower compared to cells treated with JQ1 (Wilcox test: *p* = 5.8x10^-5^) or largazole (Wilcox test: *p* < 2.22x10^-16^) (Figure 2C) alone. Noted that not all SE elements are totally susceptible to the inhibitory effect of JQ1 resulting in only partial eviction of BRD4. Such is the case of the SE encompassing the Myc gene (Figure 2A, right superenhancer). Remarkably, the combinatorial drug approach using JQ1 and largazole is sufficient to compensate for the limited inhibitory effect of single BRD4 inhibitor treatment on superenhancers like the one near MYC (Figure 2A). Again, our data suggest that largazole and JQ1 can displace BRD4 independently and mechanism of displacement may not totally overlap.

**Figure 2.**
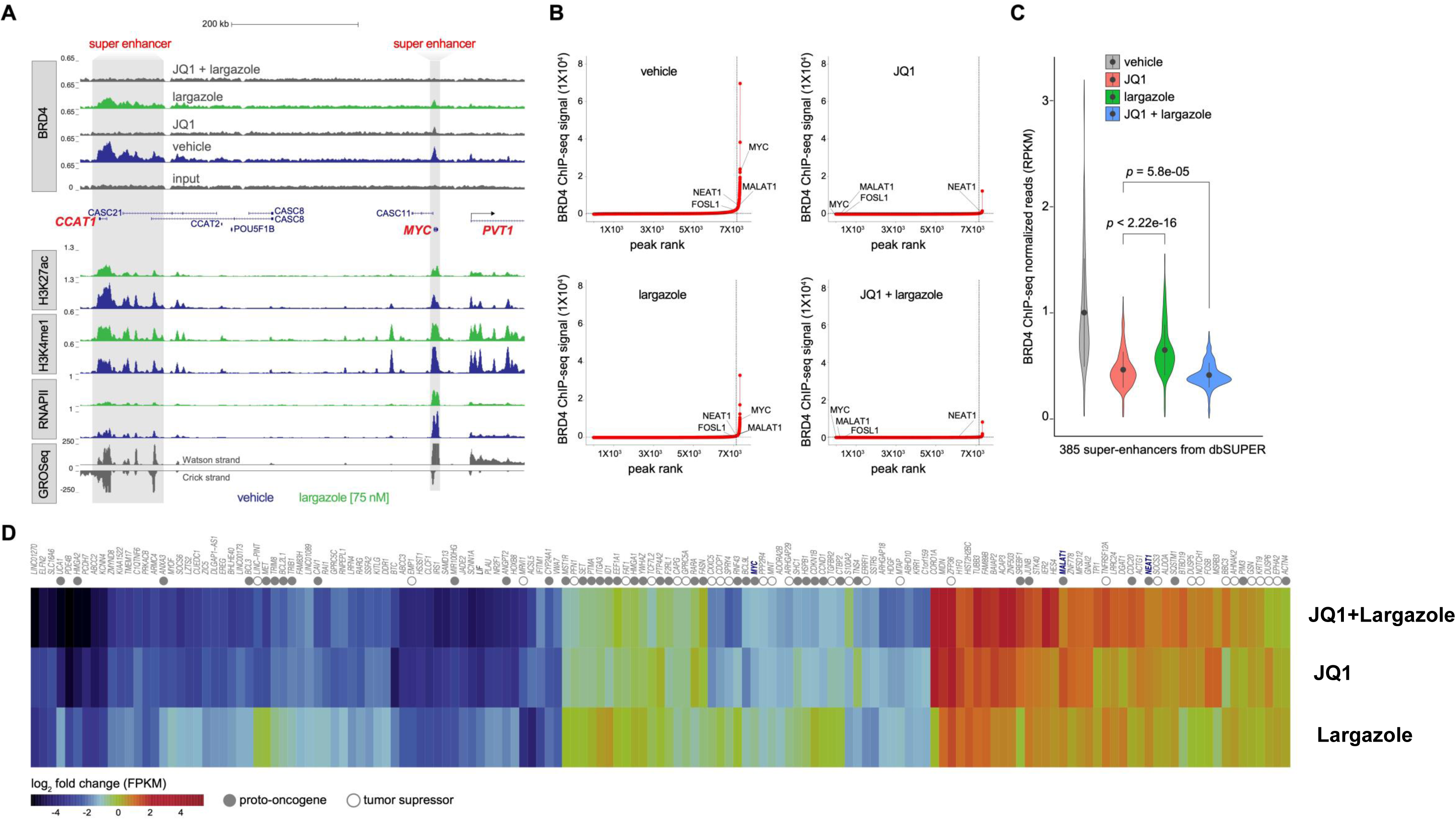
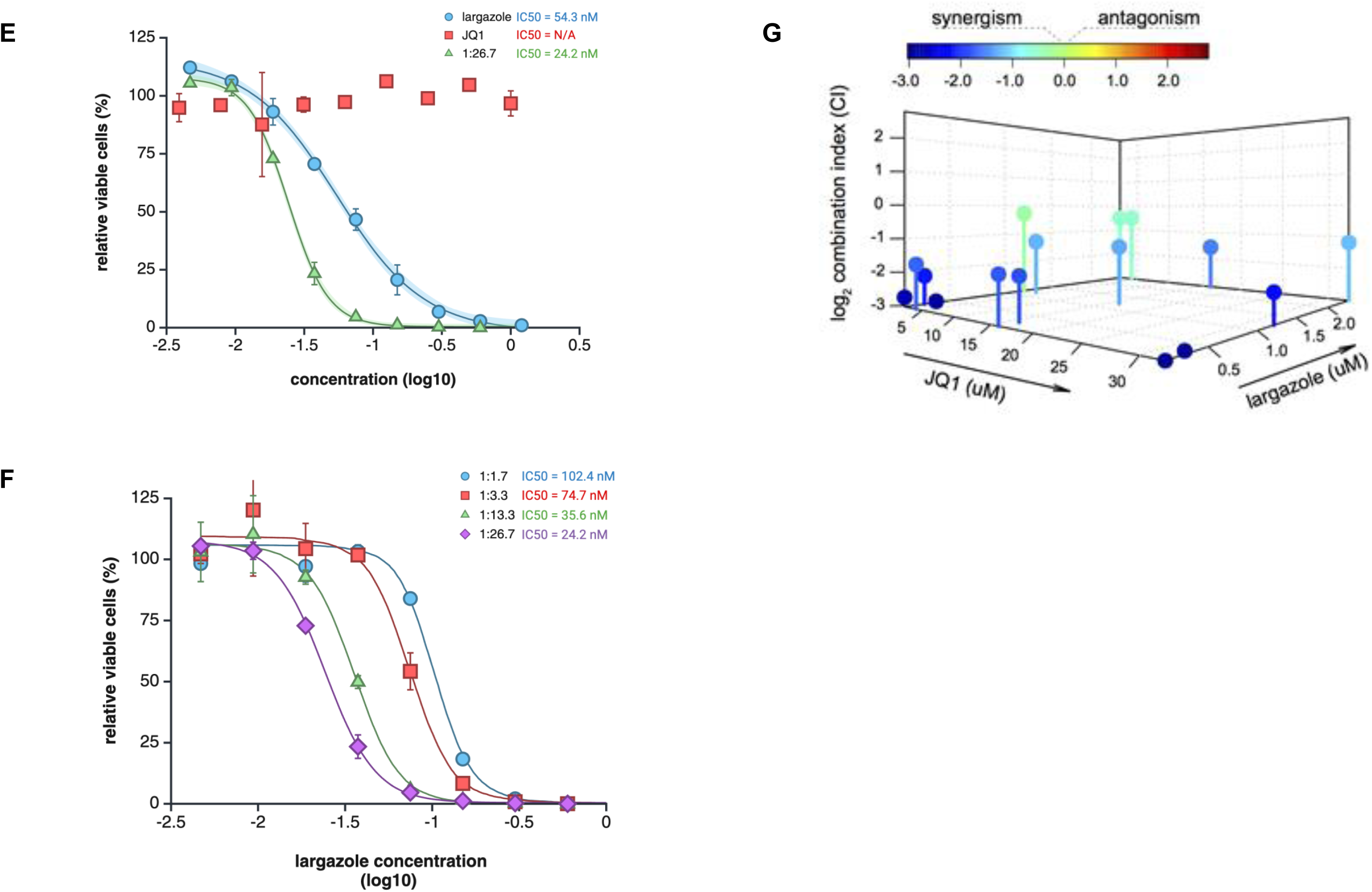
Largazole plus JQ1 treatment disrupts super-enhancers and amplifies expression changes of SE-associated genes. (**A**) Snapshot of BRD4 ChIP-seq peaks under various inhibitor treatments (vehicle control [DMSO], 75 nM largazole, 10 μM JQ1, or both inhibitors combined) in HCT116 cells after 16 hours. Shaded regions highlight two super-enhancers. (**B**) Total BRD4 signal in units of reads per million in superenhancer regions under four treatment conditions using the ROSE algorithm. (**C**) Violin plot showing BRD4 ChIP-seq signal distribution at 385 super-enhancers (dbSUPER) in HCT116 cells. The read count within each super-enhancer boundary is normalized to both super-enhancer length (kilobases) and the total number of mapped reads. (**D**) Heat map illustrating changes in mRNA accumulation from super-enhancer-associated genes. Fil led circles represent proto-oncogenes, while empty circles indicate tumor suppressor genes. (**E**) Synergistic dose-dependent growth curves of HCT116 cells treated with largazole and JQ1 at a 1:26.7 ratio to test for synergy. IC_50_ values for each condition were derived by curving fitting. 95% confidence intervals were shown in light shade with the same color for each curve. (**F**) Two-dimensional growth curves measuring synergy in HCT116 cells, where largazole and JQ1 were combined in the indicated ratios. IC50 values were derived by curve fitting. (**G**) Assessment of synergy between largazole and JQ1 using the Chou–Talalay method. Combination indices (CI) are shown in a heat map, indicating synergistic, additive, or antagonistic effects in HCT116 cells under the indicated conditions.

Next, we determined whether transcription of SE-associated genes are affected by largazole, JQ1 or combined treatment of both. A transcriptome analysis at a matching time point (16 hr) illustrates that largazole, JQ1 or the two together decrease the accumulation of SE-associated transcripts significantly (Figure 2D). Notably, the combined treatment results in an additive or synergistic effect on the accumulation changes of most SE-regulated transcripts, as observed by the augmented changes of mRNA levels with a list of representative genes (Figure 2D). Largazole treatment results in a decrease in SE activity, along with a commensurate drop in BRD4 occupancy at these sites.

Because HDAC and BRD4 inhibitors are being developed as anticancer drugs, we determined their effects on cell proliferation alone or in combination. While largazole is a potent inhibitor of HCT116 as expected (IC_50_=54.3 nM in this study), surprisingly, JQ1 treatment alone has little if any growth inhibitory activity in this cell line (EC_50_> 20 µM) (Figure 2E). When JQ1 was mixed with largazole at various ratios, we found that increasing JQ1 ratio in the drug mixture resulted in enhanced potency of largazole (Figure 2F). Using the Chou-Talalay method and Calcusyn drug combination experimental design, we observed that JQ1 can potentiate growth inhibitory activity of largazole when they are combined showing synergistic activity in a range of combined treatment conditions (Figure 2G). This result suggests that displacement of BRD4 by JQ1 in HCT116 is insufficient to induce growth inhibition. It also underscores that largazole has additional activities in transcriptional regulation and cell growth inhibition beyond displacing BRD4 from the chromatin. There has to be at least one additional regulator to be disrupted by largazole treatment.

### HDAC inhibition by largazole decreases chromatin accessibility of a subset of SEs

Since SEs identified in various cell types are located at chromatin-accessible regions ^16^, we performed quantitative transposase-accessible chromatin with sequencing (ATAC-Seq) ^17^ by incorporating Drosophila chromatin throughout sample processing as an internal reference to enable cross treatment comparisons (Figure 3). As expected, largazole treatment significantly increases the ATAC signal, which is consistent with enhanced chromatin accessibility due to hyperacetylation of histones. About 19,112 ATAC peaks show significant increase (*padjust*<0.05) compared to vehicle treated (Figure 3A). Interestingly, about 1088 regions of chromatin were found to exhibit decreased accessibility. Most strikingly, the reduced ATAC signal is higher at SEs (red dots) (Figure 3A). Using the well-established epigenetic marks, epigenetic readers, and nascent RNA as inputs into ChromHMM annotation bioinformatics tool ^18,19^, we segmented the chromatin of HCT116 cells at the basal state (vehicle treated) into 11 different states/functional regions annotated with published datasets (Figure 3B left panel). We subsequently mapped ATAC-seq reads to these bins (Figure 3B). As shown in Figure 3B, the chromatin regions that show more opening upon largazole treatment are domain insulators, poised enhancers and genic enhancers. On the other hand, active promoter regions defined by (H3K36m3, H3K27ac, H3K4me3, nascent RNA and BRD4) and some active enhancers show significant decrease in ATAC-seq peaks upon largazole exposure (Figure 3B right panel). For example, increased ATAC-seq reads was observed in the CDKN1A regulatory region with largazole (Figure 3C) while reduced ATAC-seq peaks were seen in the SE region near CASC21/CASC8 with the same treatment (Figure 3D). These results suggest that largazole reduces chromatin accessibility of a subset of SEs while increases general accessibility of chromatin.

**Figure 3.**
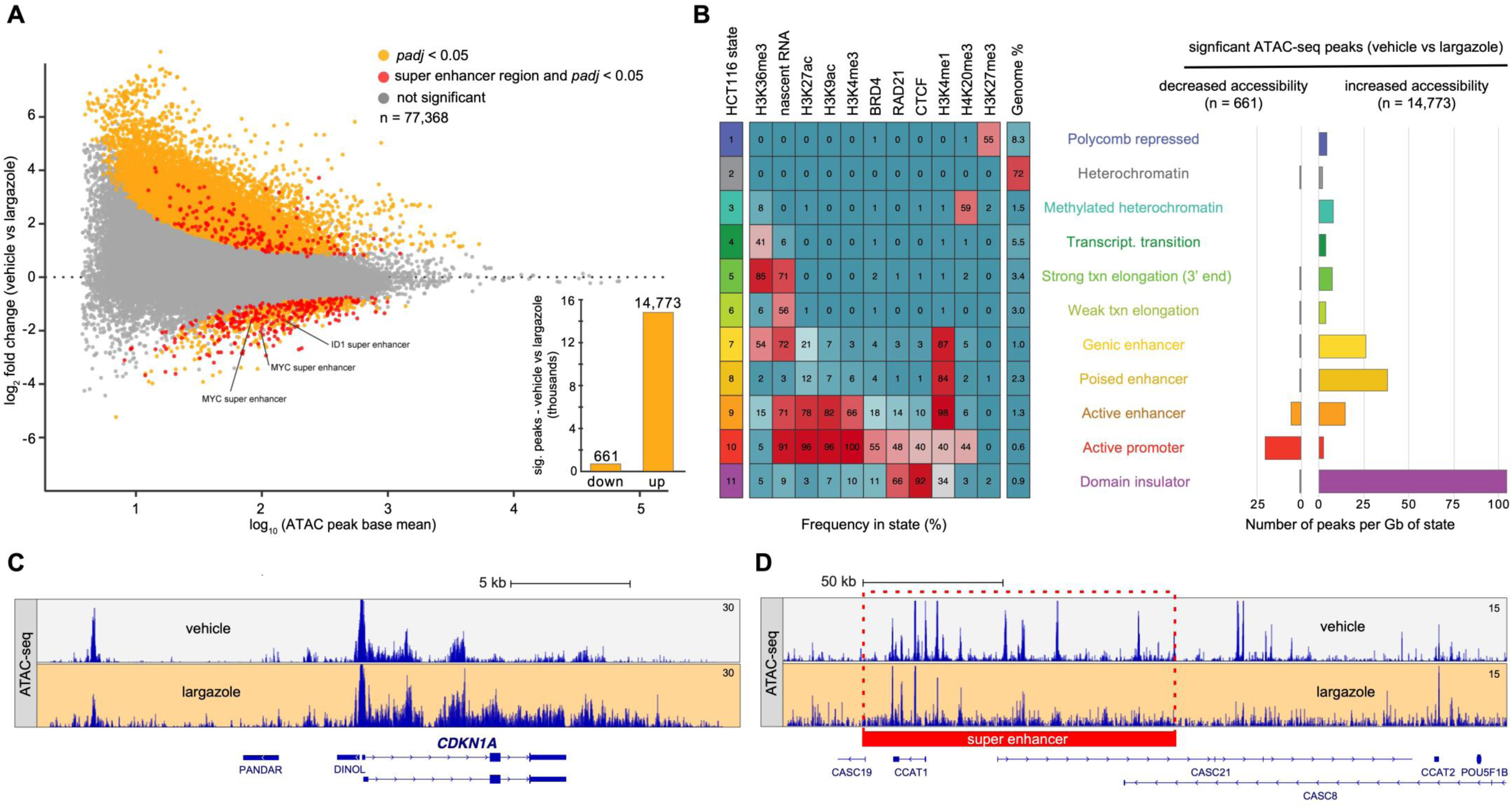
Largazole reduces chromatin accessibility at SEs and discrete proximal promoters while increasing chromatin accessibility at insulating domains. (**A**) MA plot showing changes in chromatin accessibility following 12 h of treatment with 100 nM largazole. Genomic sites exhibiting statistically significant changes (*padjust* < 0.05), determined through DESeq2 analysis ^56^, are highlighted in orange, with those located within super-enhancer regions indicated in red. (**B**) ChromHMM heatmap ^19^ depicting an 11-state chromatin roadmap of HCT116 cells, generated using data from nascent RNA, seven histone modifications (H3K36me3, H3K27ac, H3K9ac, H3K4me3, H3K4me1, H4K20me3, and H3K27me3), and three chromatin interactors (BRD4, RAD21, and CTCF). The intensity of the red color reflects the probability of observing the respective mark in each state. Candidate-state descriptions and corresponding color codes are shown to the right. The adjacent bar graph illustrates the genomic distribution of ATAC-seq peaks significantly altered by largazole: peaks with reduced accessibility appear on the left, and those with increased accessibility appear on the right. (**C**, **D**) Genome browser snapshots of the CDKN1A locus (**C**) and the most upstream MYC super-enhancer (**D**) depicting ATAC-seq signal in HCT116 cells treated with either 100 nM largazole or vehicle (DMSO) for 12 h.

### Transcription factor enrichment analysis (TFEA) reveals depletion or enrichment of key transcription factor binding sites by largazole

Since largazole causes a profound changes in chromatin accessibility and by extension likely alters transcription factor (TF) binding at their cognate motifs, we wondered which TF binding site accessibility was dramatically impacted by largazole treatment and could be responsible for SE suppression and largazole-induced growth inhibition in cancer cells. To explore this, we conducted TFEA analysis, a motif enrichment method that calculate TF motif enrichment taking consideration of both position of the motif from a designated point of interest and relative changes in sequencing reads ^20,21^. TFEA ranks accessible chromatin regions, defined by ATAC peaks, using a ranking metric based on *p*-values and the directionality of signal changes. It scans for TF motifs within the regions, and calculates the E-Score, representing the percentage enrichment or depletion of a TF motif in the ATAC regions. For each transcription factor of interest, the E-Score is determined by constructing an enrichment curve based on motif positions relative to the centers of ATAC peaks and quantifying the area between this curve and the random expectation (the diagonal). The statistical significance of each E-Score is empirically calculated by shuffling the rank order of the ATAC regions and recomputing a simulated E-Score 1000 times. We plotted the corrected E-Score for each TF motif and the number of ATAC peaks where that motif is found for 680 transcription factors with well-established binding sites defined by ChIP-seq studies ^22^ (Figure 4A). Several transcription factors exhibited significant enrichment or depletion between the control and treated conditions, with those having *p*-values<1e-35 labeled in red. We also generated plots (Figure 4B) depicting chromatin accessible regions ranked by their differential changes and *p*-values. Figure 4B provides examples of binding sites for SP1, YY1, and CTCF within these ranked accessible regions. Notably, we identified 5428 binding motifs associated with SP1 in accessible regions, with a corrected E-Score of -0.201. This indicates an enrichment of SP1 binding sites in regions of chromatin that lost accessibility upon largazole treatment. To further validate these TFEA predictions, we mapped 661 largazole-repressed and 14,773 largazole-increased accessible regions to the published ChIP-seq dataset of SP1, YY1, and CTCF in HCT116 ^23,24^. We observed a clear enrichment of SP1 and YY1 sites in the decreased regions i.e., diminished chromatin accessibility, while CTCF sites were highly enriched in the increased chromatin accessible regions (Figure 4B), which is in agreement with previous ATAC-seq results in cutaneous T cell lymphoma ^25^. In summary, these findings strongly suggest that transcription factors such as SP1 and YY1 are prime candidates responsible for mediating downstream biological effects of the HDAC inhibitor largazole.

**Figure 4.**
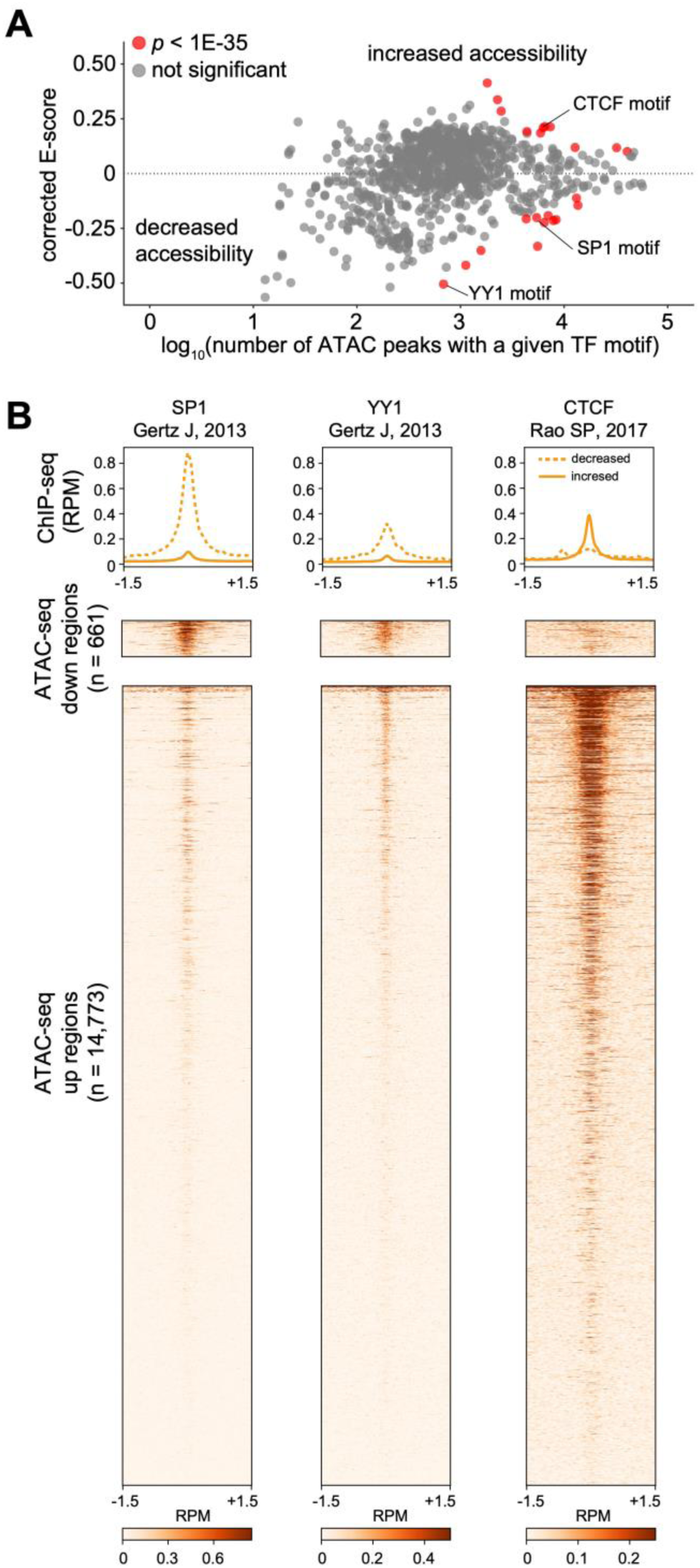
Largazole modulates chromatin accessibility at genomic sites occupied by SP1, YY1, and CTCF. **(A)** Transcription Factor Enrichment Analysis (TFEA) (ref) based on changes in ATAC-seq accessibility from vehicle and largazole-treated HCT116 cells predicts specific TFs activity. TFs showing significant alterations are highlighted in red. The MA plot displays TFs predicted to be associated with increased accessibility (e.g., CTCF) or decreased accessibility (e.g., SP1) following a 12-hour treatment with 100 nM largazole in HCT116 cells. (**B**) ChIP-seq read counts (Reads Per Million-mapped) for SP1, YY1, and CTCF are illustrated across ATAC-seq peaks that are significantly altered by largazole. Each peak is centered at the ATAC-seq peak summit (defined by MACS2) and spans 1.5 kb to each direction.

### Largazole-mediated HDAC Inhibition reduces chromatin accessibility at a subset of SEs bound by SP1 and suppresses expression of SP1-dependent genes

TFEA analysis suggests that there is a statistically significant co-localization of SP1 binding motifs with sites that lose accessibility (Figure 4), we hypothesized that SP1 binding to some of these sites is suppressed in largazole-treated cells. To test this hypothesis, we conducted SP1 ChIP-seq studies using both vehicle and largazole-treated cells. As shown in Figure 5A, among the peaks that are identified as significant SP1 binding sites (*padj*<0.05), approximately 548 exhibited reduced intensity compared to the control, while only 65 sites displayed increased intensity. Notably, when we separated SP1 sites associated with super-enhancers (highlighted in red) from those that are not, most of SP1 sites linked to super-enhancers shows decreased intensity upon largazole treatment. This result suggests that SP1 is removed from chromatin, particularly, in the SE regions.

**Figure 5.**
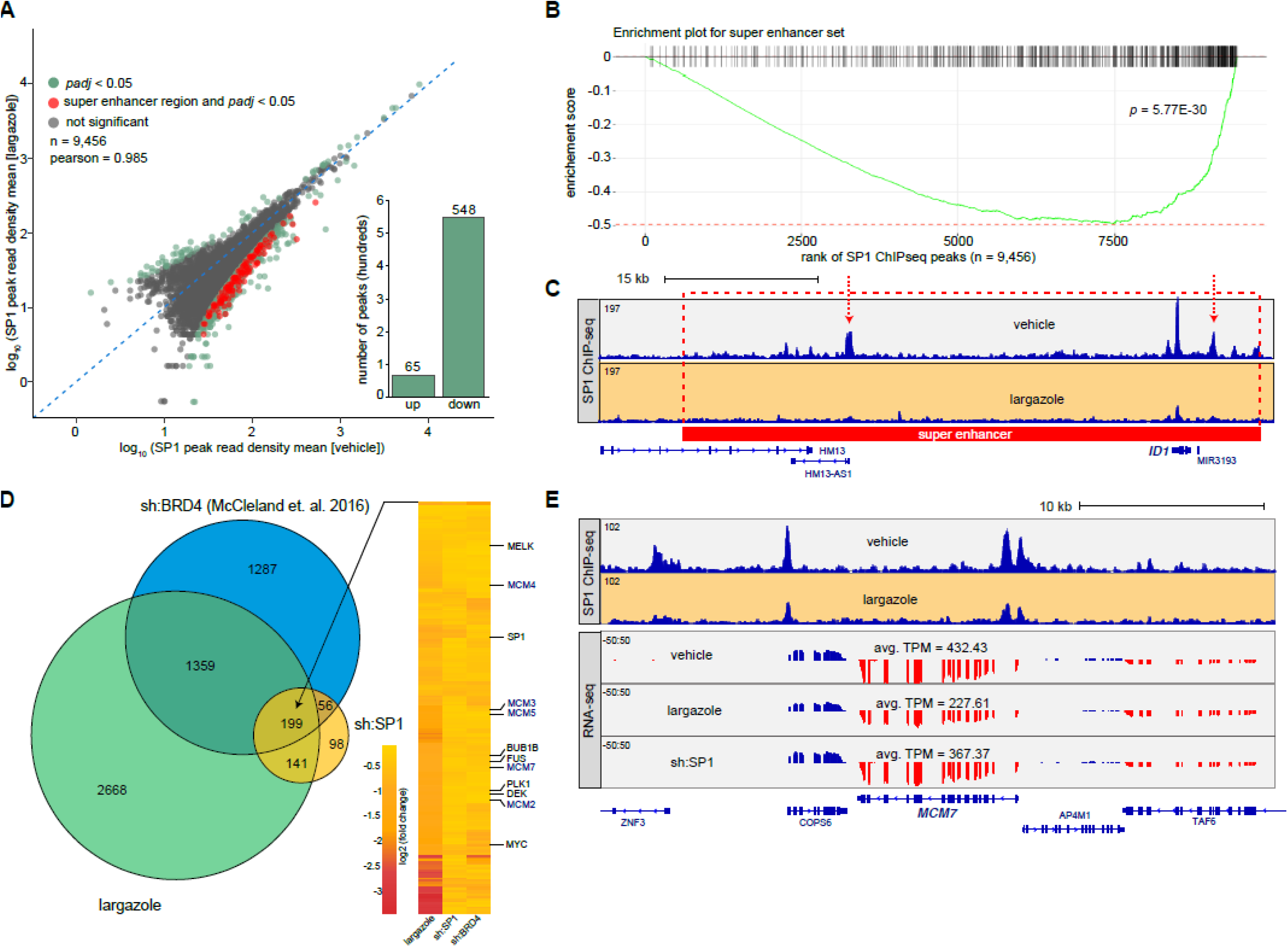
Largazole displaces SP1 from super-enhancer regions and downregulates a mitotic gene set co-regulated by SP1 and BRD4. **(A)** Scatter plot visualizing changes to SP1 ChIP-seq genomic accumulation following a 12 h largazole treatment. Genomic sites exhibiting significant changes in SP1 binding (padj < 0.05), as determined by DESeq2 analysis, are depicted in green and in red when the SP1 peak resides within super-enhancer regions (n = 113). Bar graph illustrating the effect of largazole on SP1 ChIP-seq peaks undergoing higher (n = 65) or lower (n = 548) occupancy. (B) Enrichment plot for the set of SP1 ChIP-seq peaks localized within super enhancer regions (n = 498) illustrated as vertical black lines (total SP1 ChIP-seq peaks = 9,456). **(C)** Visualization of the ID1 locus in genome browser displays SP1 ChIP-seq signals in HCT116 cells treated with 100 nM largazole or vehicle control (DMSO) for 12 hours. Bounderies of the ID1-super enhancer are illustrated by a red-dotted line and the significant differentially occupied SP1 ChIP-seq peaks are denoted as black dotted rectangles. **(D)** Venn diagram comparing significant differentially downregulated transcripts from HCT116 cells treated with either 12h largazole [100 nM], shRNA-mediated downregulation of SP1, or shRNA-mediated downregulation of BRD4 ^53^. The heatmap illustrates the shared gene targets for the three conditions (n = 199) and highlights a mitotic gene battery. **(E)** Genome browser visualization of the MCM7 gene locus illustrating SP1 ChIP-seq (top) and RNA-seq (bottom) signals in HCT116 cells treated with 100 nM largazole or vehicle control (DMSO).

To further determine if the SP1 ChIP-seq datasets from vehicle and largazole treated cells show statistically significant differences in SE binding upon drug treatment, we performed fGSEA (Fast Gene Set Enrichment Analysis) ^26^ with ranked SP1 ChIP-seq peaks by log-fold change. This analysis identified significant reduction of SP1 peaks associated with SEs in largazole treated samples, as indicated by high negative ES score for the SE dataset (*p*<5.77e-30) and FDR<0.05) (Figure 5B), suggesting depleting of SP1 from SEs potentially contributing to largazole drug responses. Figure 5C shows an example of SP1 eviction from the SE surrounding ID1, a gene that is known to promote tumor growth, metastasis and resistance to drug and radiation therapy ^27^. Depletion of SP1 at SEs aligns with our previous observation of inactivation of SEs by largazole and other HDIs, providing a basis for further investigation into the antitumor effects of this class of drugs.

Since SP1 is widely known as transcriptional activator, loss of SP1 is expected to result in altering the expression of genes it regulates. To identify SP1-dependent genes in HCT116, we depleted SP1 with an siRNA and determined the changes in transcriptomic expression by RNA-seq analysis. Functional enrichment analysis with GSEA revealed that SP1 is heavily involved in RNA processing, DNA replication and DNA repair process (Supplemental Figure 3). Next, we investigated whether we can identify overlapping genes that were downregulated across largazole, SP1 depletion (siRNA) or BRD4 depletion (shRNA) treatments. The overlap analysis identified 199 shared downregulated genes and the heatmap illustrates the changes of these genes across three treatments (Figure 5D). These overlapping genes are involved in DNA replication (e.g. MCMs), mitotic progression and spindle checkpoint regulation (e.g. BUB1B, MELK, DEK and PLK1). Figure 5E shows an example of correlation between reduction in SP1 occupancy seen in ChIP-seq data and downregulation of MCM7 transcripts under conditions to largazole treatment or SP1 depletion. These results suggest that largazole treatment can cause the loss of SP1 and BRD4 in super-enhancers, potentially contributing to the observed transcriptional repression in largazole-treated cells.

### Largazole suppresses SP1 expression in tumor cells and depletion of SP1 sensitizes largazole-induced growth inhibition

The absence of SP1 on chromatin could be attributed to either SP1 redistribution or the depletion of intracellular SP1. To investigate which mechanism is likely responsible for the absence of SP1 on chromatin, we conducted RNA-seq studies on HCT116 cells exposed to increasing doses of largazole. As depicted in Figure 6A, the RNA-seq data reveal a dose-dependent decrease in SP1 mRNA levels as largazole doses increase. This dose-dependent reduction in SP1 mRNA was independently confirmed through RT-qPCR analysis, as shown in Figure 6B. Next, we assessed SP1 protein levels in cells treated with largazole over a 48-hour period. Consistent with the suppression of SP1 mRNA expression, SP1 protein levels displayed a time-dependent depletion in HCT116 cells (Figure 6C). Since SP1 is a member of the SP1 family of transcription factors, which comprises a group of proteins known for their ability to bind to specific DNA sequences and regulate various genes, we aimed to investigate the specificity of largazole-induced SP1 suppression. To address this question, we measured the levels of SP4, another member of the SP1 family that shares structural similarities with SP1, using Western blot analysis. Unlike SP1, SP4 remains unchanged following largazole treatment, suggesting that largazole-induced reduction in SP1 levels is specific (Figure 6D). Additionally, we determined whether the effect of largazoe on SP1 is cell line specific. As shown in Figure 6D and 6E, largazole induced SP1 reduction was similarly observed in the human pancreatic tumor cell line MiaPaCa and the mouse head and neck squamous carcinoma cell line TCh3. These results suggest that largazole induced SP1 suppression is specific and occurs in multiple cell lines.

**Figure 6.**
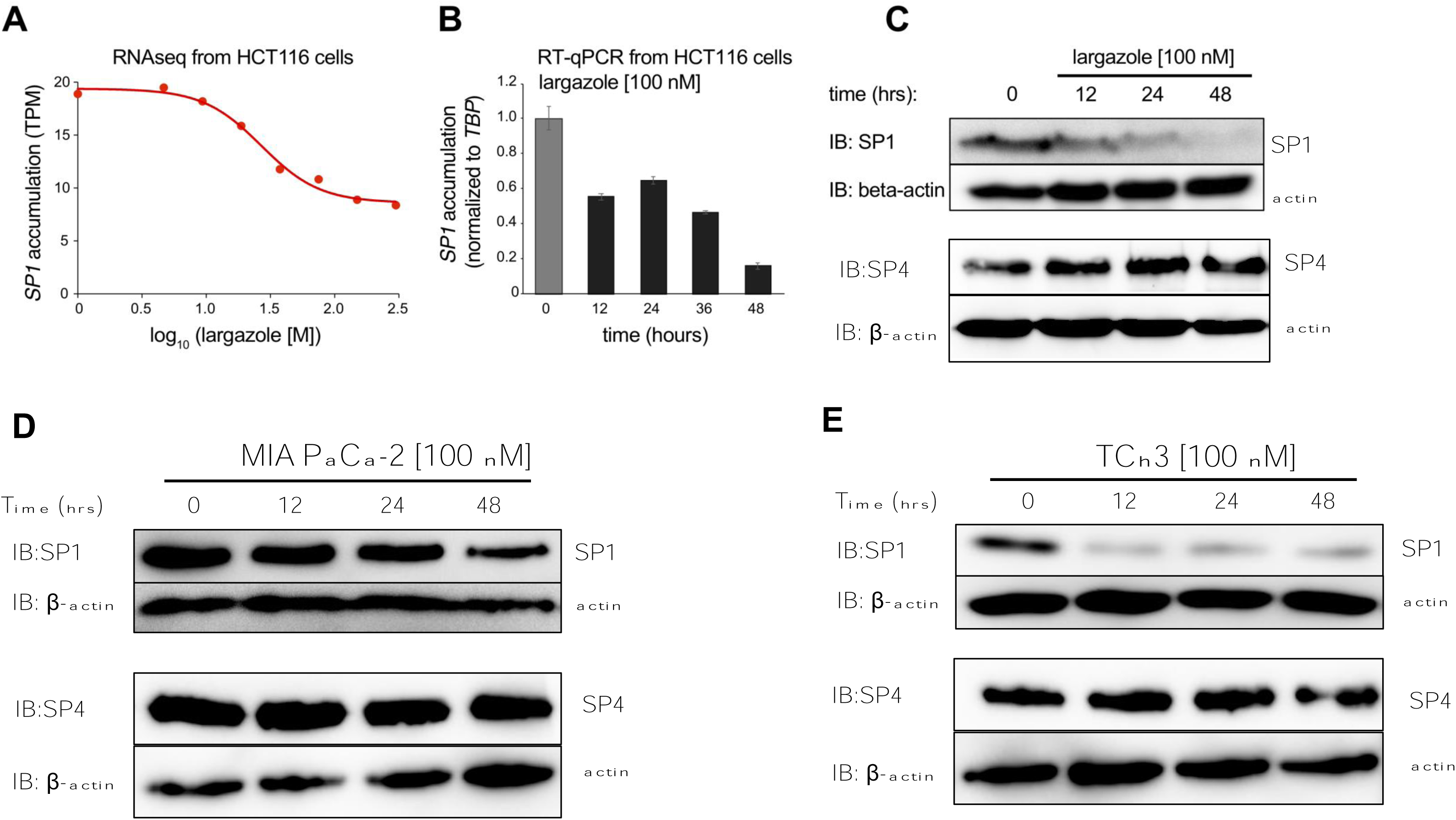
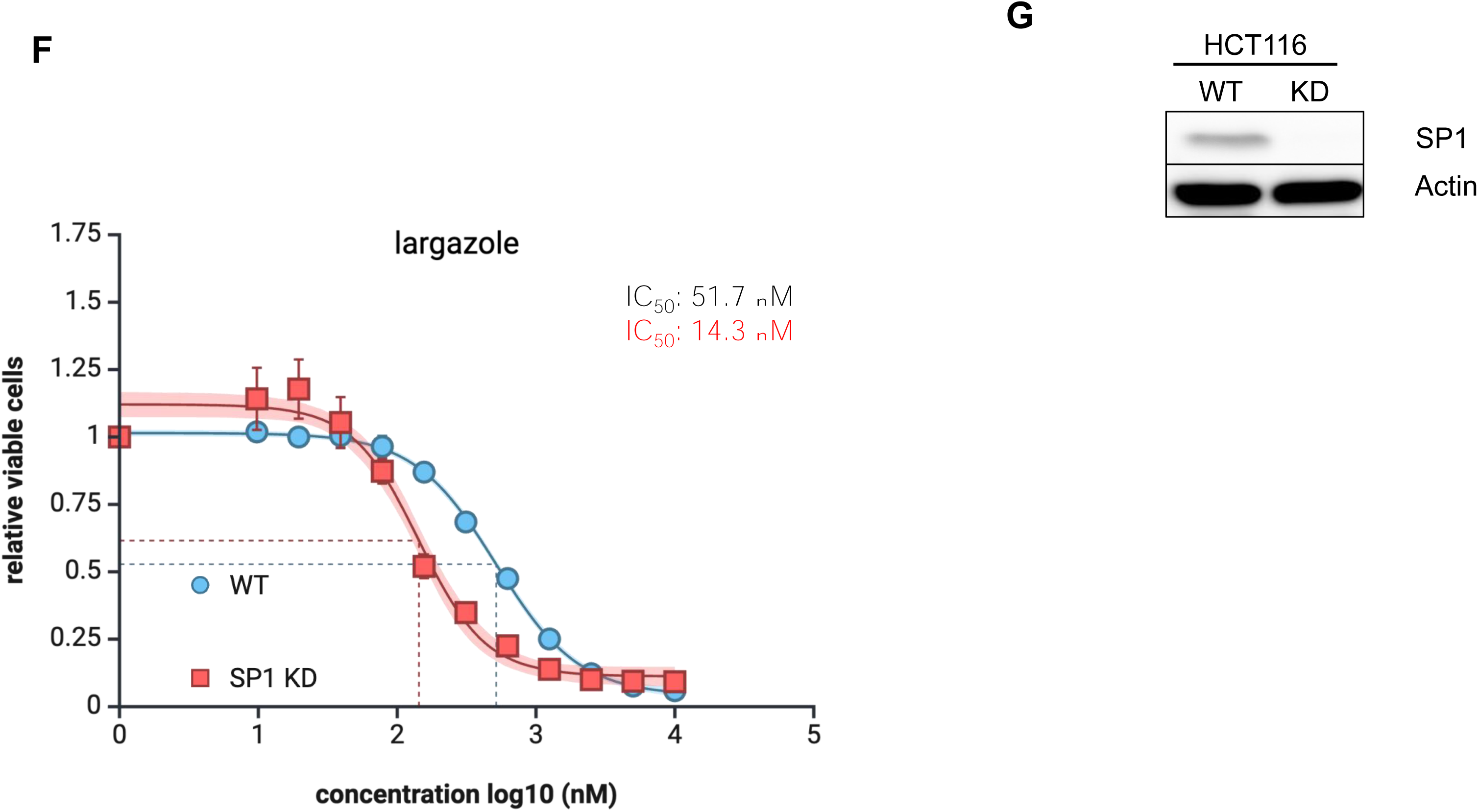
Largazole suppresses SP1 mRNA and protein expression and depletion of SP1 sensitizes largazole-induced growth inhibition. (**A**) Total RNA was harvested from HCT116 cells treated with DMSO, 9.4 nM, 18.8 nM, 30 nM, 37.5 nM, 75 nM, 150 nM and 300 nM for 16 hrs and subjected to RNA-seq analysis. SP1 transcript levels measured by TPM (transcript per million) at each dose of largazole treatment were plotted. (**B**) Relative SP1 transcripts levels in HCT116 cells treated with 100 nM largrazole for indicated times (hrs) were measured by RT-qPCR analysis and plotted relative to vehicle treated samples. Data are represented as mean ± SD, n=3. Western blot analysis of SP1 and SP4 protein levels treated with 100 nM largazole for indicated times in HCT116 MiaPaca (**D**) and mouse squamous carcinoma cell line TCh3 (**E**). β-actin was blotted as the loading control.(**F**) Dose-dependent growth curves of HCT116 and HCT116 SP1 KD, a stable cell line expressing SP1 shRNA. Largazole IC_50_ values for the two cell lines were derived from curving fitting. (**G**) Western blotting of SP1 in cell lysates from wild type SP1 and SP1 KD.

To investigate whether the suppression of SP1 is involved in largazole-induced tumor cell growth inhibition, we depleted SP1 in HCT116 using shRNA. We then evaluated the impact of SP1 depletion on largazole-induced growth inhibition. Notably, the IC_50_ of largazole for HCT116 cells stably expressing SP1 shRNA (∼14.3 nM) was approximately 3.6-fold lower than that of wild-type cells (∼51.7 nM), as presented in Figure 6F and Figure 6G. This underscores that reducing SP1 expression significantly enhances the sensitivity of HCT116 cells to largazole. These findings suggest that largazole primarily inhibits SP1 expression across multiple cell lines through the suppression of mRNA expression. The down-regulation of SP1 may represent a key mechanism by which largazole inhibits cell growth.

### Largazole-induced SP1 depletion prolongs prometaphase and metaphase during HCT116 mitosis

Previously we found that largazole induces dose-dependent cell cycle arrest leading to the accumulation of 4N cells ^5^. The findings that largazole disrupts the architecture of SEs by suppression of SP1, BRD4 and Cohesins prompt us to investigate whether perturbations of SEs by largazole are linked to its activity to regulate cell cycle progression. To delve deeper in to the molecular mechanisms behind largazole-induced cell cycle arrest, we conducted live imaging of HCT116 cells exposed to largazole. This revealed that a significant portion of treated cells exhibited defects in chromosome alignment and failed mitosis (Figure 7A). Building upon these findings, we conducted an in-depth investigation into the impact of largazole on chromosome alignment and mitotic progression. In brief, we synchronized HCT116 cells through a double thymidine block and then released them into nocodazole to induce mitotic arrest. Subsequently, cells arrested in mitosis were released from the G2/M phase by removing nocodazole and adding largazole at indicated concentrations, along with MG132, which inhibits the metaphase-to-anaphase transition. Chromosome alignment was assessed by evaluating chromosome alignment in fixed cells. Quantitative analysis of the effect of largazole on chromosome alignment revealed that increasing largazole exposure led to significant defects in chromosome alignment, with defective alignment observed in up to 50% of cells treated with 500 nM largazole. Higher concentrations of largazole resulted in mitotic failure and an increased rate of mitotic catastrophe (Figure 7B). To further characterize the mitotic defects induced by largazole treatment, we conducted live cell imaging analysis and quantified mitotic progression in the presence or absence of largazole. In vehicle-treated cells, the duration of prometaphase and the time required for complete chromosome alignment to metaphase were approximately 20 minutes and 30 minutes, respectively (Figure 7C, top panel). However, in the presence of largazole, a significant fraction of cells experienced mitotic failure, the extent of which depended on the largazole dose (Figure 7C, middle and bottom panels). Cells that entered mitosis displayed either prolonged prometaphase (>150 minutes) or an extended time to achieve metaphase chromosome alignment (>175 minutes). These results suggest that largazole disrupts mitotic progression by interfering with chromosome alignment or inducing mitotic failure.

**Figure 7.**
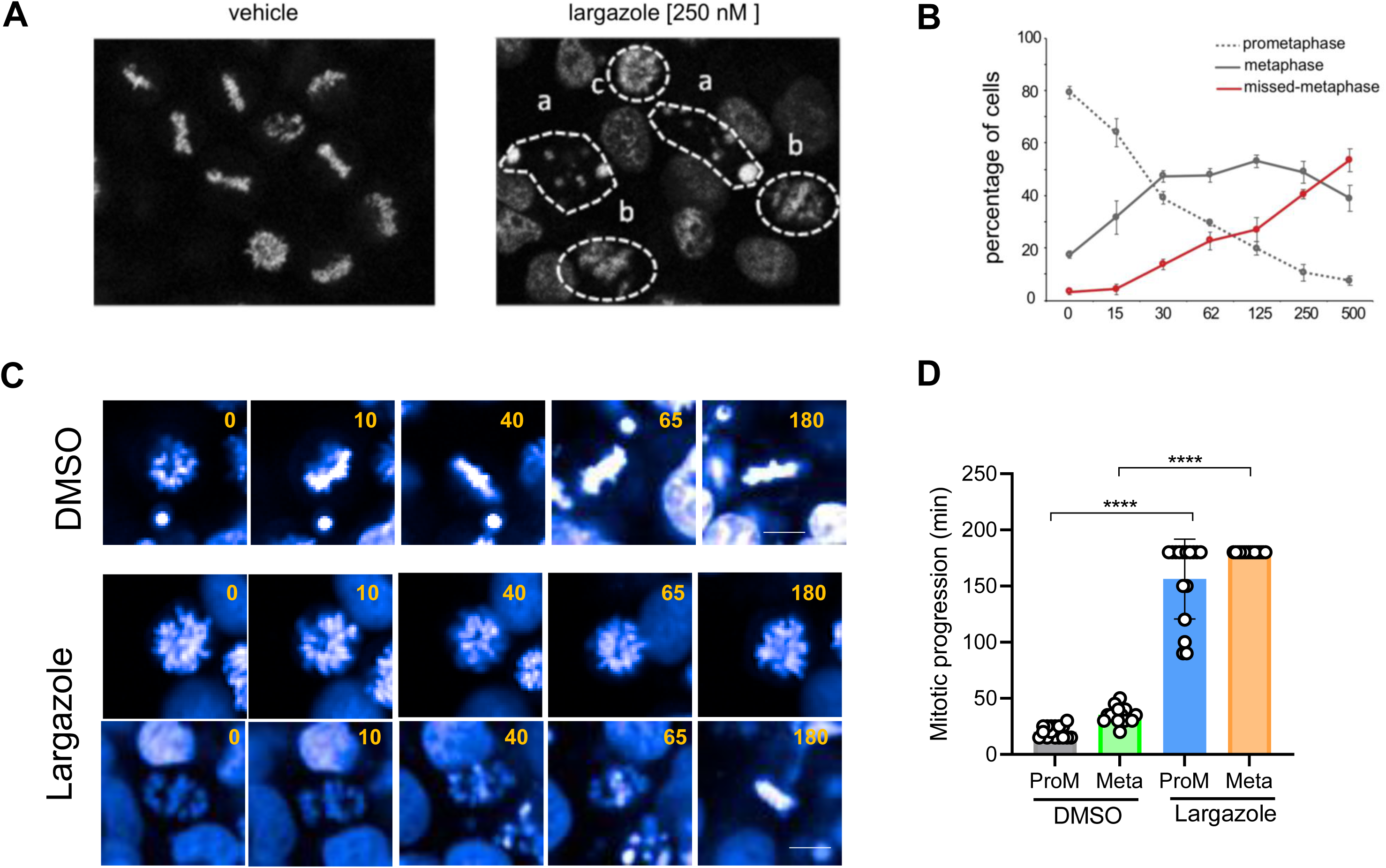
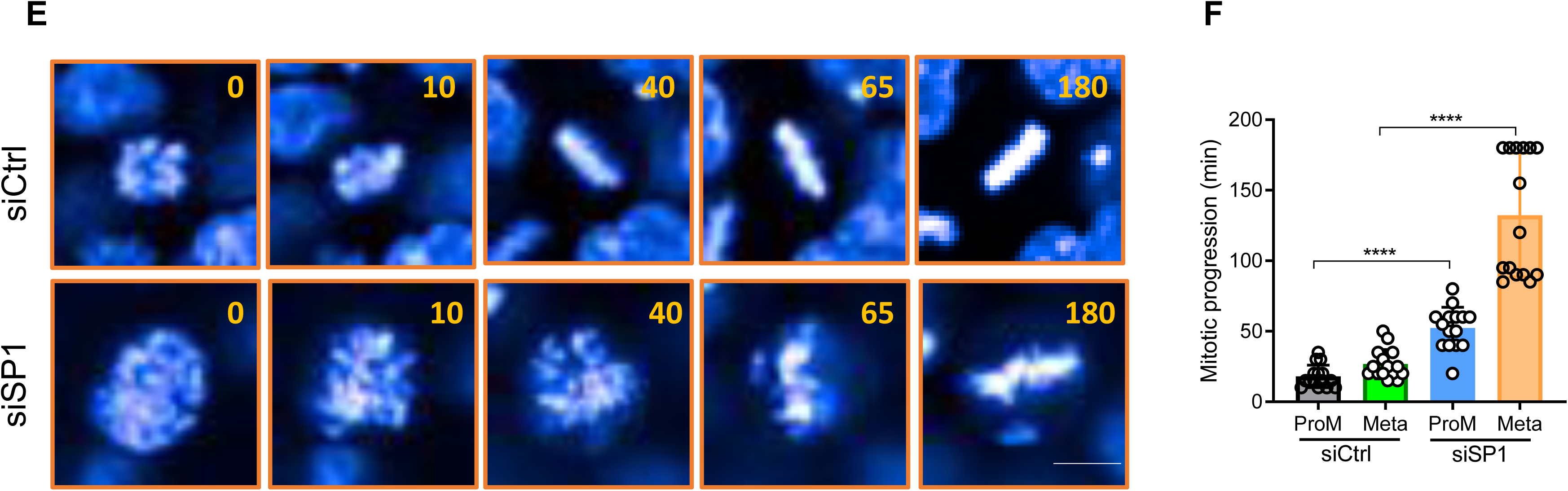
Largazole induces chromosome mis-alignment and delays mitotic progression. HCT116 cells were treated 2 μM thymidine for 24 hours. After 24 hours, cells were washed with PBS 3 times, and treated with 10 μM RO-3306 with DMSO or various concentrations of largazole for 19 hours. After 19 hours, cells were washed with PBS 3 times and imaged in DMEM in the presence of 10 μM MG132 and 20 μM Hoechst 33258 for 3 hours at 10 min intervals. (**A**) Representative images of prometaphase cells treated with vehicle or 250 nM largazole. Chromosome mis-alignment or delayed alignment is seen with the largazole treatment (dashed circles). (**B**) Quantitation of largazole effect on mitotic progression. X-axis is the concentration of largazole used. Y-axis, represents the percentage of cells in prometaphase, metaphase or missed-metaphase in largazole treated cells. (**C**) Representative montages of live cell imaging of vehicle and largazole treated cells during mitosis. (**D**) Quantitation of duration of the length of prometaphase and metaphase in control and largazole treated cells. Total 18 cells were tracked. Y axis shows the duration of prometaphase (ProM) or times for achieving a full alignment of chromosomes (Meta); the duration longer than 180 min were recorded as 180 min. Data are represented as mean ± SD. Student’s t-test was used for statistical analysis for data. \**p < 0.05*, \*\**p < 0.01*, \*\*\**p < 0.001*, \*\*\*\**p < 0.0001*. (**E**) Representative montages of live cell imaging of control and SP1 shRNA treated cells during mitosis. (**F**) Quantitation of duration of the length of prometaphase and metaphase in control and SP1 shRNA HCT116 cells. Total 17 cells were tracked manually. Y axis shows the duration of prometaphase (ProM) or times for achieving a full alignment of chromosomes (Meta); the duration longer than 180 min were recorded as 180 min. Data are represented as mean ± SD. Student’s t-test was used for statistical analysis for data. *\*p < 0.05, **p < 0.01, ***p < 0.001, ****p < 0.0001*.

To determine whether largazole-induced SP1 depletion and SE deactivation contribute to defects in chromosome alignment and delayed mitotic progression, we examined the effects of SP1 depletion using HCT116 SP1 knockdown cells. Both wild-type and mutant cells were synchronized as described earlier. We observed that SP1 knockdown cells exhibited notable defects in chromosome alignment and took significantly longer to enter prometaphase or achieve chromosome alignment to complete metaphase. The mitotic phenotypes of SP1 knockdown cells resembled those of largazole-treated cells (Figure 7C and Figure 7D). These results suggest that largazole induces defects in chromosome alignment and mitotic progression, and such defects can be attributed to the loss of SP1 caused by largazole treatment. This underscores largazole-induced SP1 suppression as one of the primary underlying mechanisms contributing to the antiproliferative activity of largazole.

## DISCUSSION

Here, we investigated the molecular mechanisms by which largazole, an HDAC inhibitor, suppresses gene expression and inhibits cancer cell growth. Our findings demonstrate that largazole treatment predominantly increases global chromatin accessibility, but it also leads to decreased chromatin accessibility in specific genomic regions, particularly a subset of super-enhancers (SEs). Through bioinformatics analysis and ChIP-seq studies, we discovered that largazole treatment displaces both SP1 and BRD4 proteins from SEs, resulting in the disruption of SE architecture. Additionally, we showed that largazole suppresses the expression of SP1 mRNA, leading to disturbances in chromosome alignment during mitosis and ultimately causing mitotic failure due to the suppression of SP1 and cohesin. These results provide insights into how the HDAC inhibitor largazole suppresses gene expression by modulating chromatin accessibility and introduce a novel anti-cancer mechanism involving the disruption of mitotic progression and erasing transcription memories.

HDAC inhibitors are known to increase histone acetylation, leading to a more open chromatin structure and increased accessibility of transcription factors and RNA polymerase to gene promoters. Consequently, HDI treatment often results in the upregulation of previously repressed genes. However, it has been observed that HDIs can also downregulate many genes in a dose-dependent manner ^3,5,28,29^. The mechanisms behind HDAC inhibitor-induced gene repression are complex and not fully understood. Some studies suggest that an increase in RNA polymerase II (RNAPII) pausing may contribute to transcriptional repression for certain genes, as HDAC enzymes are required for transcription elongation or the release of paused RNAPII ^4,5,30,31^. We and others have noted that HDIs selectively target a subset of SEs and their associated transcripts for suppression ^5,32–36^. One potential explanation for the inactivation of SEs under HDAC inhibitor treatment is the disruption of normal contacts between SEs and gene bodies, which can compromise the fidelity of SE-gene pairing ^33^. BRD4, a protein with two bromodomains that bind to acetylated histone tails, is thought to facilitate contacts between SEs and promoters through a looping mechanism ^14,37^. Since HDAC inhibitor treatment elevates overall histone acetylation in the genome and causes acetylation marks to spread into gene bodies ^5,32,33,38^, hyperacetylated histones may sequester BRD4, reducing the available pool of BRD4 required for mediating interactions between promoters and enhancers ^32,39^. Our data (Figure 1 and Figure 2) supports the idea that BRD4 deficiency at SEs and disrupted architecture contribute to the deactivation of SEs and the repression of genes such as c-Myc under largazole treatment. Notably, our ATAC-seq and ChIP-seq data indicate a disruption of SE architecture, including a decrease in chromatin accessibility and the loss of YY1 and SP1. (Figure 4 and Figure 5). Additionally, our western blot results (not shown) also show that largazole suppresses STAG2 and RAD21, core components of the cohesin complex known to mediate contacts between SEs and promoters ^40^. Disruption of SE architecture may lead to the disabling of positive elongation factor b (P-TEFb)-dependent RNA polymerase II pause release, further contributing to gene repression.

The observed reduction of BRD4 at SEs seems to be counterintuitive, as SEs are highly enriched with H3K27ac marks ^41^. Our ATAC-seq results shed light on this phenomenon. While largazole treatment results in greater chromatin openings overall in agreement with previous observations with other HDIs ^25,31,33^, some sites exhibit decreased chromatin accessibility, with SEs being preferentially affected. Our analysis suggests that SP1 binding sites are associated with the loss of chromatin accessibility (Figure 3), a finding confirmed by SP1 ChIP-seq analysis (Figure 4). Intriguingly, the SP1 sites that experience reduced chromatin accessibility contain a higher frequency of co-localized BRD4 sites (Figure 5F). This observation suggests a potential interaction between SP1 and BRD4 at a subset of SEs, and the loss of SP1 at SEs may be responsible for the reduction in BRD4 levels. This hypothesis is in agreement with previously finding that universal stripe factors (USFs), which comprises 30 SP, KLF, EGR, and ZBTB family members known for binding overlapping GC-rich sequences, provide chromatin accessibility for partnering factors in mammalian genome ^42^. This raises the possibility that USFs may be the gatekeeper for SEs and set the stage of SEs associated factor such as BRD4 to mediate transcriptional activation. Future experiments are needed to elucidate the relationship between SP1, BRD4, and SEs and their impact on gene regulation.

The largazole-induced reduction of SP1 binding to chromatin could result from various mechanisms, including posttranslational modifications, suppression of SP1 protein or mRNA synthesis, or increased SP1 protein degradation ^43,44^. While our data indicates that largazole suppresses SP1 mRNA synthesis, we cannot exclude the possibility of other mechanisms contributing to SP1 eviction from chromatin. Downregulation of the SP1 family of transcription factors has also been observed in in Rhabdomyosarcoma cell lines upon exposure to HDI panobinostat and vorinostat ^45^. Notably, we observed specific suppression of SP1 but not SP4, distinguishing it from the earlier study. SP1 is considered proto-oncogenic, as it is often overexpressed in various human cancers and regulate genes critical for cell proliferation and survival ^43,44^. In line with this, our GSEA analysis of SP1 knockdown compared to wild type samples revealed that SP1 regulates expression of genes that are heavily involved in DNA replication, DNA repair, RNA processing and protein translation, which aligns with the hypothesis that SP1 is a critical cell growth promoting gene. Further support this view, our experiments demonstrated that partial depletion of SP1 sensitizes cells to largazole-induced growth inhibition, suggesting that antagonizing SP1 is one of the mechanisms by which largazole inhibits tumor growth *in vitro* and *in vivo*. What remains to be determined is how largazole suppresses SP1 specifically without affecting other SP1 factors. Sp1 gene is known to be autoregulated by SP1 due to the presence of multiple SP1 binding sites at its promoter ^46^. Displacement of SP1 from its genomic sites may trigger a feedback loop to ramp down its expression. Future studies need to address whether largazole also regulates SP1 via posttranslational modifications and degradation.

Despite five HDIs being approved for cancer treatments, fundamental mechanisms of their antitumor effects on cancer remain debatable. They can induce various cellular responses, including apoptosis, cell cycle arrest, oxidative stress, DNA damage, and differentiation ^47,48^, while also affecting processes such as angiogenesis and immune responses (Wang et al. 2019; Qu et al. 2017; Rafehi et al. 2014). It is still uncertain which molecular processes disrupted by HDIs contribute the observed anticancer effects predominantly. One recurring theme is that HDIs cause global changes in the epigenetic landscape and the biological outcomes are highly context-dependent. Largazole, in particular, disrupts chromosome alignment and mitotic progression through the suppression of SP1, providing a potential mechanistic explanation for its anti-cancer effects. Previous studies have shown HDIs can induce mitotic slippage and disruption of spindle assembly checkpoint (reviewed in ^49^). Others reported that SP1 delocalizes to mitotic centromeres and rapid ablation of SP1 at mitotic onset results in chromosome segregation errors and aberrant mitotic progression ^50,51^. Our result links HDI induced chromosome mis-alignment to an early genomic response to HDIs in SP1 eviction and disruption of SEs. It is worth to note that BRD4, like SP1, also localizes on condensed chromosomes during mitosis ^49,51^. Their retentions on mitotic chromosome may be part of mitotic bookmarking required for the maintenance of transcriptional programs and chromosome accessibility into G1 ^52^. A corollary to a requirement for SP1 and BRD4 for maintaining transcription memory would be that largazole, probably other HDIs too, can reset transcriptional programs by erasing this memory after mitosis via targeting SP1 and BRD4. This proposed mechanism of action may be relevant to the observed therapeutic effects of HDIs as chromatin accessibility of tumor and T cells from responders to HDI therapy appear to swing back to the pattern of normal cells ^25^. Future studies should evaluate whether SP1 suppression is a response biomarker that could predict the response to therapy for various indications in clinical relevant samples.

### Key resources table

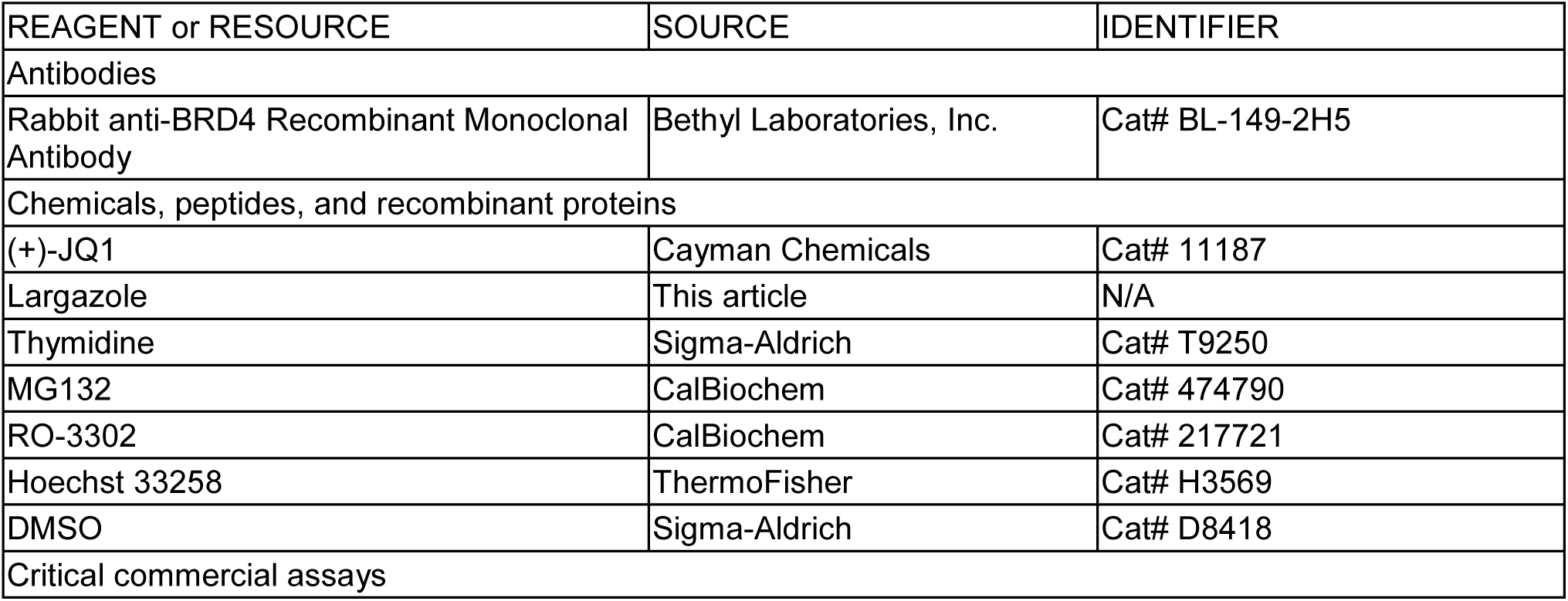
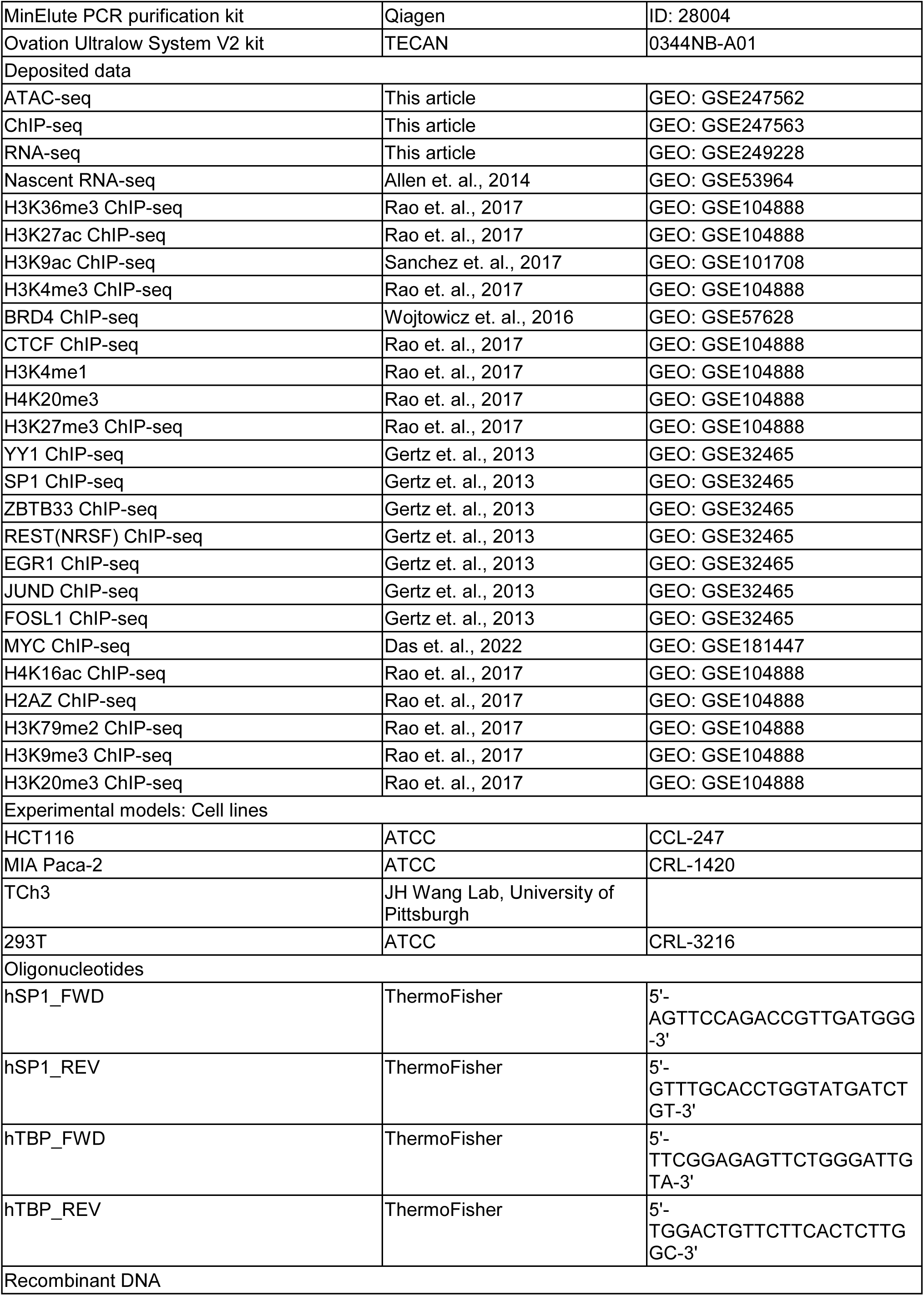
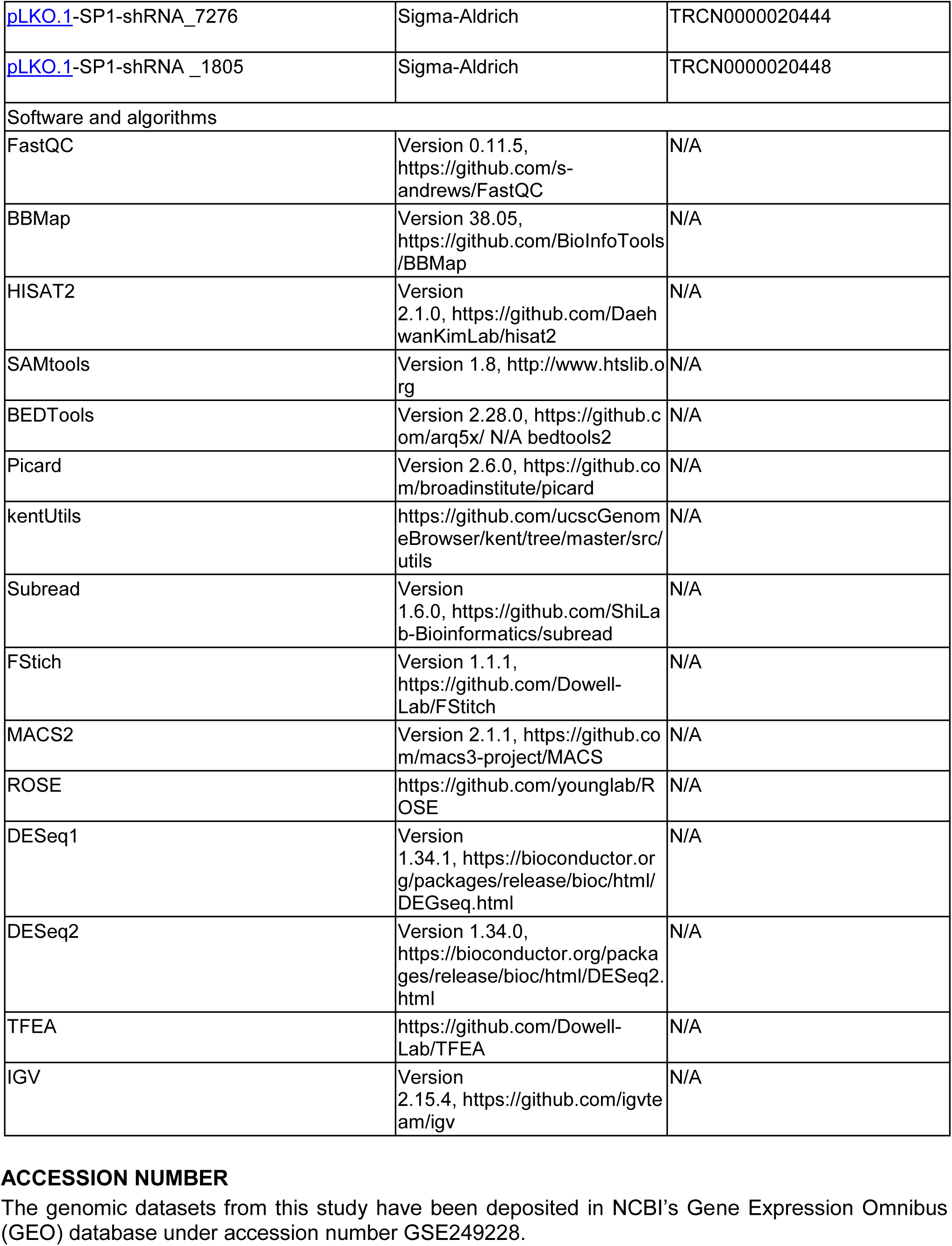

## ACCESSION NUMBER

The genomic datasets from this study have been deposited in NCBI’s Gene Expression Omnibus (GEO) database under accession number GSE249228.

## ACKNOWLEDGMENTS

We thank members of Dowell and Liu laboratories for discussion and suggestions. This work was supported by grants from National Institutes of Health R01GM144749 (Liu) and R01CA229174A1 (Wang). E.B. and G.W. were supported by a predoctoral training grant from NIGMS (T32GM08759). The ImageXpress MicroXL was supported by a NCRR grant S10RR026680, and FACSAria was supported by S10OD021601, and Opera Phenix imaging system by S10OD025072 from NIH.

## AUTHOR CONTRIBUTIONS

G.S., Z.L., Q.X., and X.L. designed the study and analyzed the data. G.S., Z.L., Q.X., G.W., and C.T. performed experimental studies. G.S., Z.L., S.H., Q.X., and X.L. wrote the manuscript in consultation with D.T., M.A., and R.D. with inputs from all authors.

## CONTACT FOR REAGENT AND RESOURCE SHARING

Additional information and requests for resources and reagents should be directed to and will be fulfilled by the Lead Contact, Dr. Xuedong Liu.

## EXPERIMENTAL MODEL AND SUBJECT DETAILS

### Cell lines and cell culture

The human colorectal carcinoma (HCT116), pancreatic carcinoma (MiaPaCa-2), and the embryonic kidney epithelial (293T) cell lines were acquired from ATCC. The murine TCh3 cell line was generated by the JH Wang Lab at the University of Pittsburgh from a female K15.CrePR1(+)p53^f/f^PIK3CA^c/c^ mouse ^54^. All cells were grown in Dubelcco’s modified Eagle medium (DMEM) supplemented with 10% fetal bovine serum (FBS), 1% penicillin-streptomycin, and 1%

GlutaMAX (Invitrogen). Cell lines were maintained at 37°C with 5% CO_2_ and tested routinely for mycoplasma contamination. For experiments with inhibitor treatments, cells were treated with either vehicle (DMSO) or the indicated inhibitor concentration for 12 h or 16 h before harvesting for transcriptome (RNA-seq) or chromatin state (ChIP-seq) analysis.

## METHOD DETAILS

### Immunoblotting

Western blots were carried out using standard protocols. Briefly, cells were grown, treated, and harvested as previously stated. Cell lysates were separated by SDS-PAGE and transferred to GVS nitrocellulose 0.22 um membranes. Blots were probed with primary antibodies, followed by peroxidase-conjugated secondary antibodies (GE Healthcare Life). Signal for all immunoblots was acquired using the ImageQuant LAS 4000 biomolecular imager (GE Healthcare LS) with an average exposure of 30 s. Antibodies used are as follows: SP1 (Santa Cruz Biotechnology, sc-17824); SP4 (Santa Cruz Biotechnology, sc-390124); RAD21 (Cell Signaling, cat. # 4321); STAG2 (Cell Signaling, cat. # 5882); beta-Actin (SC-47778).

### shRNA-Mediated Knockdown

For shRNA-mediated gene silencing, we used two SP1-specific shRNA that target starting position nt1805 or nt7276 of SP1 coding sequence (TRCN0000020448: 5’-GCTGGTGGTGATGGAATACAT-3’ and TRCN0000020444: 5’-CCCAAGTTTATTTCTCTCTTA-3’, respectively, Sigma-Aldrich). Lentiviruses were prepared by transfecting 293T cells with the desired shRNA vector along with three packaging vectors (psPAX2, pMD2.G, and VSV-G) as described. The knockdown data pertain to a polyclonal cell line acquired through lentiviral transduction and subsequent puromycin selection (1 µg/ml).

### RNA Isolation, Reverse Transcription, and qPCR

Total RNA was extracted with TRIzol LS reagent (Thermo Fisher) from HCT116 cells treated with 100 nM largazole for the indicated time points. RNA concentrations were measured using Nanodrop and 1 mg was used for cDNA synthesis using SuperScript III Reverse Transcriptase Kit (Thermo Fisher) with random hexamers. Remnant genomic DNA was eliminated with RQ1 RNase-Free DNase (Promega). cDNA was analyzed via the CFX Opus real-time PCR system (Bio-Rad Laboratories). For all experiments, relative expression levels of the target genes were determined by calculating the 2-ΔCt values. Experiments were normalized using the internal control TBP (Forward primer: 5’-CACGAACCACGGCACTGATT; Reverse primer: 5’-TTTTCTTGCTGCCAGTCTGGAC-3’) transcript levels.

### Chromatin Immunoprecipitation, Spike-In, and Sequencing

ChIP experiments were carried out as previously published. In summary, HCT116 cells underwent treatment with either vehicle or with the specified inhibitor for 12 h or 16 h, followed by cross-linking with 1% formaldehyde for 10 minutes at room temperature and immediate quenching using 250 mM glycine. After two washes with PBS, the cell membranes were disrupted using a hypotonic buffer (50 mM NaCl, 1% NP-40 alternative, 2 mM EDTA, 10 mM Tris, 1 mM DTT, 2 mM EDTA, 1X protease inhibitor cocktail (Roche # 04693124001). The nuclei were isolated by centrifugation and resuspended in lysate buffer (150 mM NaCl, 0.5% Triton X-100, 2 mM EDTA, 0.1% SDS, 20 mM Tris, 1 mM DTT, 1X protease inhibitor cocktail). Similarly, Drosophila S2 cells underwent crosslinking, followed by lysis of the cell membrane, with subsequent recovery of nuclei for spike-in controls. HCT116 resuspended samples were spiked with 3x10^5 Drosophila nuclei and sonicated for 25/35 cycles (30 s ‘on’ at high level and 30s ‘off’ per cycle) using a Bioruptor (Diagenode; Denville, NJ, USA) and spun for 10 min at 16,000 × g in a microcentrifuge. The resulting supernatant was then incubated for 5 h at 4◦C with 5– 20 µg of antibodies and 20 µl of 50% slurry with protein A beads (Millipore; Billerica, MA, USA). The immunoprecipitated chromatin was then recovered and DNA purified using phenol-chloroform extraction. Antibodies used are as follows: BRD4 (Bethyl Laboratories, Inc. cat # BL-149-2H5); SP1 (Santa Cruz Biotechnology, cat # sc-390124). ChIP samples were sequenced on the HiSeq 2500 (Illumina) at the University of Colorado Shared Resources Genomics Core Facility under the PE150 protocol.

### ChIP-seq Analysis

ChIP-seq datasets were aligned using HISAT2 mapping software v2.1.0. Reads were mapped to the hg38 reference human genome. We used SAMtools v1.3.1 to generate a sorted pileup format of the aligned reads. For each experiment, genome coverage bed graph files were generated using BEDTools v2.28.0 and then normalized by the total number of mapped reads. Normalized bed graph files were subsequently converted to bigwig files and uploaded to IGV v2.15.4 for visualization.

### RNA Sequencing and Analysis

RNA was isolated from HCT116 cells treated for 16 h using TRIzol reagent (Life Technologies) following the manufacturer’s instructions. The concentration of each sample was determined using the QubitTM 3.0 Fluorometer (Thermo Fisher), and its integrity was assessed using an Agilent Bioanalyzer 2100 (Agilent Technologies). The Illumina TruSeq RNA Sample Preparation kit was utilized to create RNA sequencing libraries. The resulting library fragment lengths were confirmed using an Agilent Bioanalyzer 2100. Library quantification was performed using the QubitTM 3.0 Fluorometer, and sequencing was conducted at the Next-Generation Sequencing Facility at the University of Colorado BioFrontiers Institute using an Illumina HiSeq 2000 sequencing system. All sequencing libraries underwent multiplexing before sequencing. Reads were pruned using BBDUK from the BBMap suite v38.05 and mapped to hg38 human genome using HISAT2 v2.1.0. After mapping, alignment files were processed using SAMtools v1.3.1. FeatureCounts from the Subread software package v1.6.0 was used to count the total number of sequencing reads that aligned to each putative gene model. To determine which genes were differentially expressed, we used the R package DESeq version 1.34.1. A subset of RNA-sequencing studies including HCT116 SP1 shRNA and WT comparison were performed at LC Sciences (Houston). RNA was isolated from vehicle or drug treated cells using PureLink RNA Mini Kit (Thermo Fisher) and submitted to the vendor for sequencing studies.

### ATAC-seq Nuclei Extraction

ATAC-seq was performed essentially as described ^17^, except nuclei were extracted before transposition. We generated biological duplicates for each ATAC-seq condition. In brief, for each condition, 100,000 cells were treated for 12h. The cells were washed with 5mL D-PBS, trypsinized, and transferred to a 50mL tube containing 5mL fresh DMEM on ice. The cells were then centrifuged at 500xg for 5min at 4°C and DMEM was replaced with ice-cold D-PBS. The pellet was resuspended in 5mL of ice-cold swelling buffer (10mM Tris-HCl pH 7.5, 2mM MgCl2, 3mM CaCl2) and incubated for 5min on ice. The cells were centrifuged at 400xg for 5min at 4°C; the supernatant was removed. The pellets were then resuspended in 5mL swelling buffer with 20% glycerol and kept on ice. Next, 5mL of lysis buffer (swelling buffer + 20% glycerol + 1% NP-40) was added gently and incubated for 5min on ice. Following this, 10mL of lysis buffer was added and centrifuged at 600xg for 5min at 4°C. Supernatant was discarded, the pellet was washed with 5mL swelling buffer + 20% glycerol, and centrifuged at 600xg for 5min at 4°C. Supernatant was discarded and the pellet was resuspended in 1mL freezing buffer (40% glycerol, 10mM Tris-HCl pH 8, 2mM MgCl2, 0.1mM EDTA), transferred to 2mL cryovial, and stored at -80°C.

### ATAC-seq Spike-in, Library Preparation, and Sequencing

Cryopreserved nuclei were sent to Active Motif for ATAC-seq. Nuclei were thawed at 37°C and washed with cold PBS. 2x10^5 HTC116 nuclei were mixed with 4x10^4 drosophila S2 nuclei per reaction. ATAC-seq was performed as previously described with some modifications ^17,55^. Briefly, spike-in nuclei were resuspended in lysis buffer, pelleted, and tagmented with Tn5 transposase at 37°C for 30min. Tagmented DNA was purified using the MinElute PCR purification kit (Qiagen), amplified for 11 PCR cycles, and purified using Agencourt AMPure SPRI beads (Beckman Coulter). Libraries were quantified with the KAPA Library Quantification Kit (KAPA Biosystems) and sequenced with PE42 sequencing on a NovaSeq 6000 sequencer (Illumina). Tagmentation, library construction and sequencing were performed by Active Motif Inc (Carlsbad, CA).

### ATAC-seq Analysis

We generated duplicate ATAC-seq libraries for each condition examined and sequenced each to a depth of ∼40 million 42-bp paired-end reads. We first pruned the reads using BBDUK from the BBMap suite v38.05 with flags: forcetrimleft=15, forcetrimright2=3, qtrim=10. Alignment of the paired-end DNA sequencing reads was performed using HISAT2 v2.1.0 to the hg38 human or the DM6 genomes with the following parameters: --very-sensitive, --no-spliced-alignment, --no-mixed, --no-discordant. To determine regions of chromatin accessibility, we employed MACS2 v2.1.1 with flags: -nomodel --shift -100 --extsize 200 --broad -B --keep-dup auto. Peak regions with a *p*-value < 1E-7 were selected for downstream analysis. Tabulation of ATAC-seq reads over the accessible regions was performed with FeatureCounts from the Subread software package v1.6.0. Differential peak enrichment analysis was performed using DESeq2 v1.34.0 by estimating size factors with the Drosophila ATAC-seq counts as the control genes. Fragment coverage was normalized by adjusting the bedgraph files to the estimated size factors previously described.

### Model Fitting of ChromHMM

For functional segmentation of the human chromatin with ChromHMM, we employed publicly available ChIP-seq and nascent RNA-seq resulting from HCT116 cells under basal conditions ^18,19^. The NGS data was processed as mentioned above, binarized as previously described, and ChromHMM was fitted with default parameters.

### TFEA Analysis

Transcription factor enrichment analysis from ATAC-seq and SP1 ChIP-seq data was conducted with the use of TFEA and employing the dataset “best_curated_Human.meme” encompassing 1279 human DNA motifs with a FIMO threshold of 1e-6. Genomic sites with read signal above background were selected as mentioned above.

### Cell Viability Assay

Cell viability of HCT116 cells, subjected to either vehicle (DMSO) or the specified largazole concentrations was assessed using crystal violet staining. Briefly, cells were rinsed once with phosphate-buffered saline (PBS) and fixed for 20 minutes at room temperature with 4% paraformaldehyde while gently rocking. Following a single PBS wash, fixed cells were stained with 0.5% crystal violet solution (Sigma) in 20% methanol for 10 minutes at room temperature. Subsequently, cells were thoroughly washed with water and air-dried overnight. Finally, 150 μl of a developing solution (methanol, ethanol, and water in a 4:1:1 ratio) was added to each well, and absorbance was measured at 560 nm. For drug combination studies, we used alamarBlue assays for measuring cell viability. Briefly, vehicle or drug treated cells growing on 96-well plate were incubated with 1x stock solution (dilute from 250x 1% Resazurin Sodium salt solution in PBS) at 37 C for >3 hr after media removal. The fluorescence intensity of each well was measured on a fluorescence spectrophotometer using 560/590 nm (excitation/emission) filter settings (SpectroMax, Molecular Devices).

### Mitotic Progression and Chromosome Alignment Analysis

HCT116 or derivative cells were synchronized by incubating them with 2 mM thymidine for 24 hours. After synchronization, cells were washed three times with D-PBS and further incubated with 10 µM RO-3306 for 19 hours to arrest them at the G2/M boundary. Following the RO-3306 treatment, cells were washed three times with D-PBS and then incubated with 10 µM MG132 and 2 µM Hoechst 33342. During this incubation, cells were imaged continuously for 3 hours using the ImageXpress XL microscope (Molecular Devices) to monitor mitotic progression and chromosome alignment.

---

**Supplemental Figure 1.**
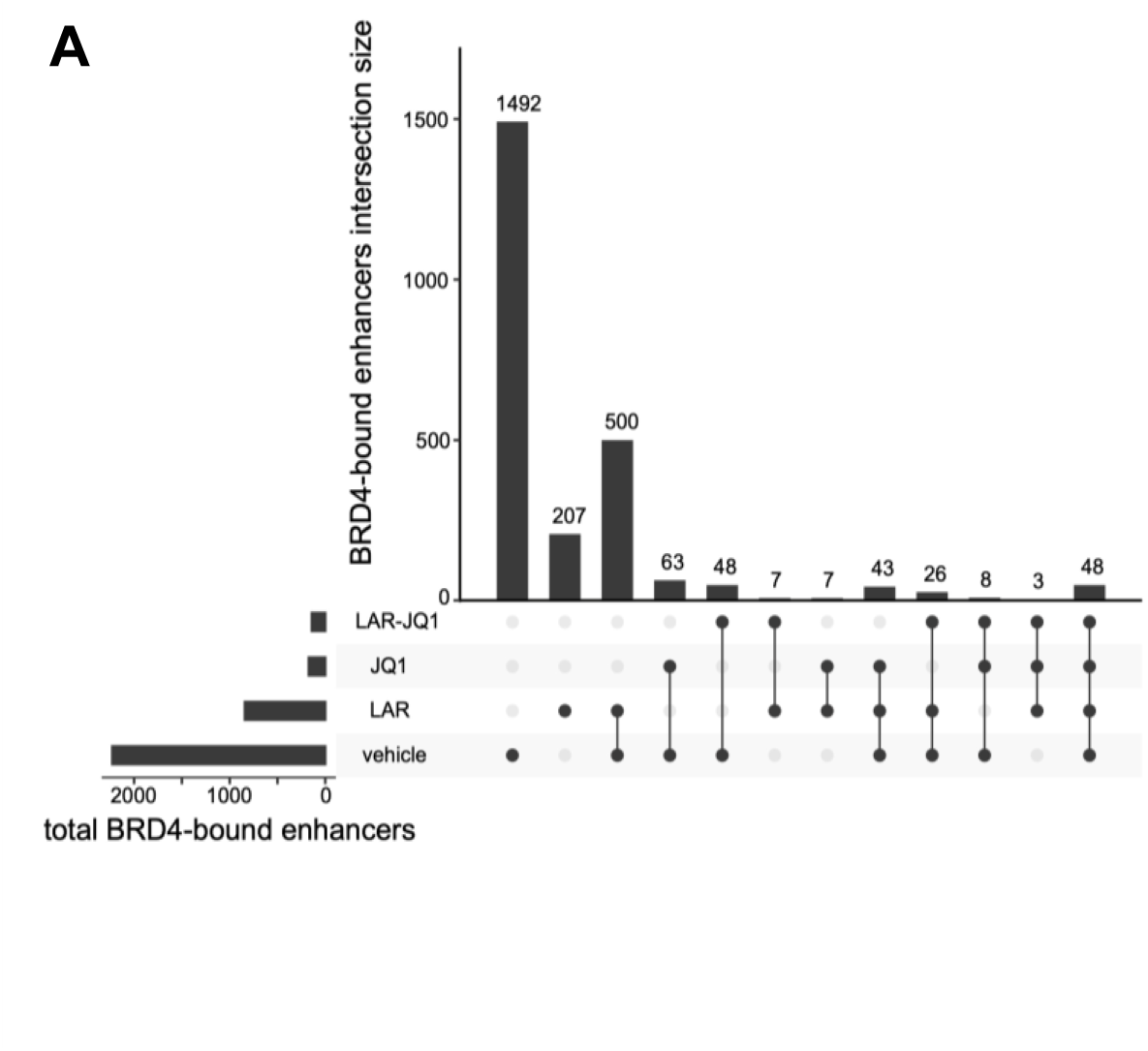
(**A**) UpSet plot illustrating the overlapping accumulation of BRD4 along 2,452 enhancer elements across four cellular treatments.

**Supplemental Figure 2.**
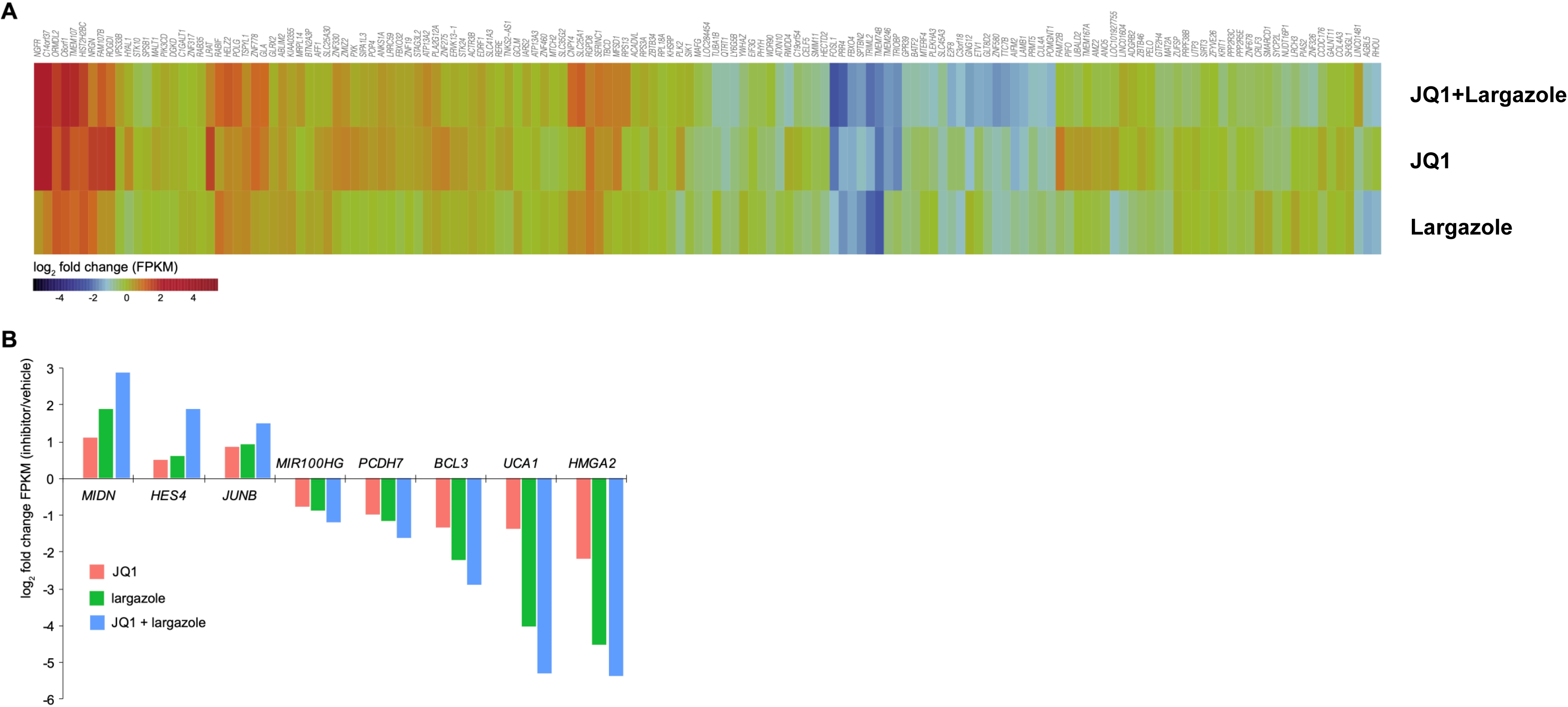
Largazole and JQ1 cotreatment amplify expression patterns in SE-associated genes. (**A**) Heat map visualization of mRNA accumulation changes for a randomly selected gene set. (**B**) Bar chart depicting steady-state mRNA levels of selected super-enhancer associated genes across four cellular conditions.

**Supplemental Figure 3.**
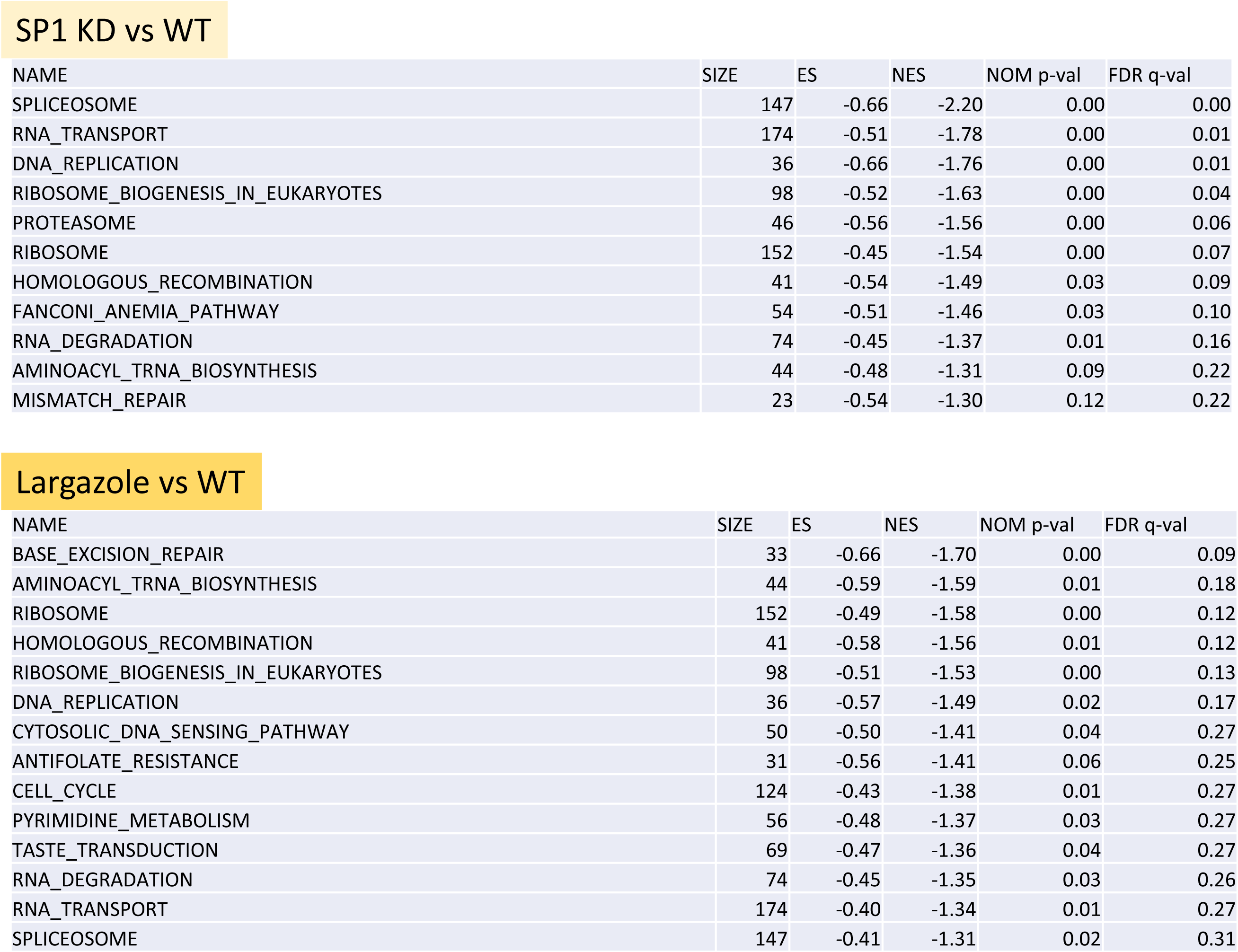
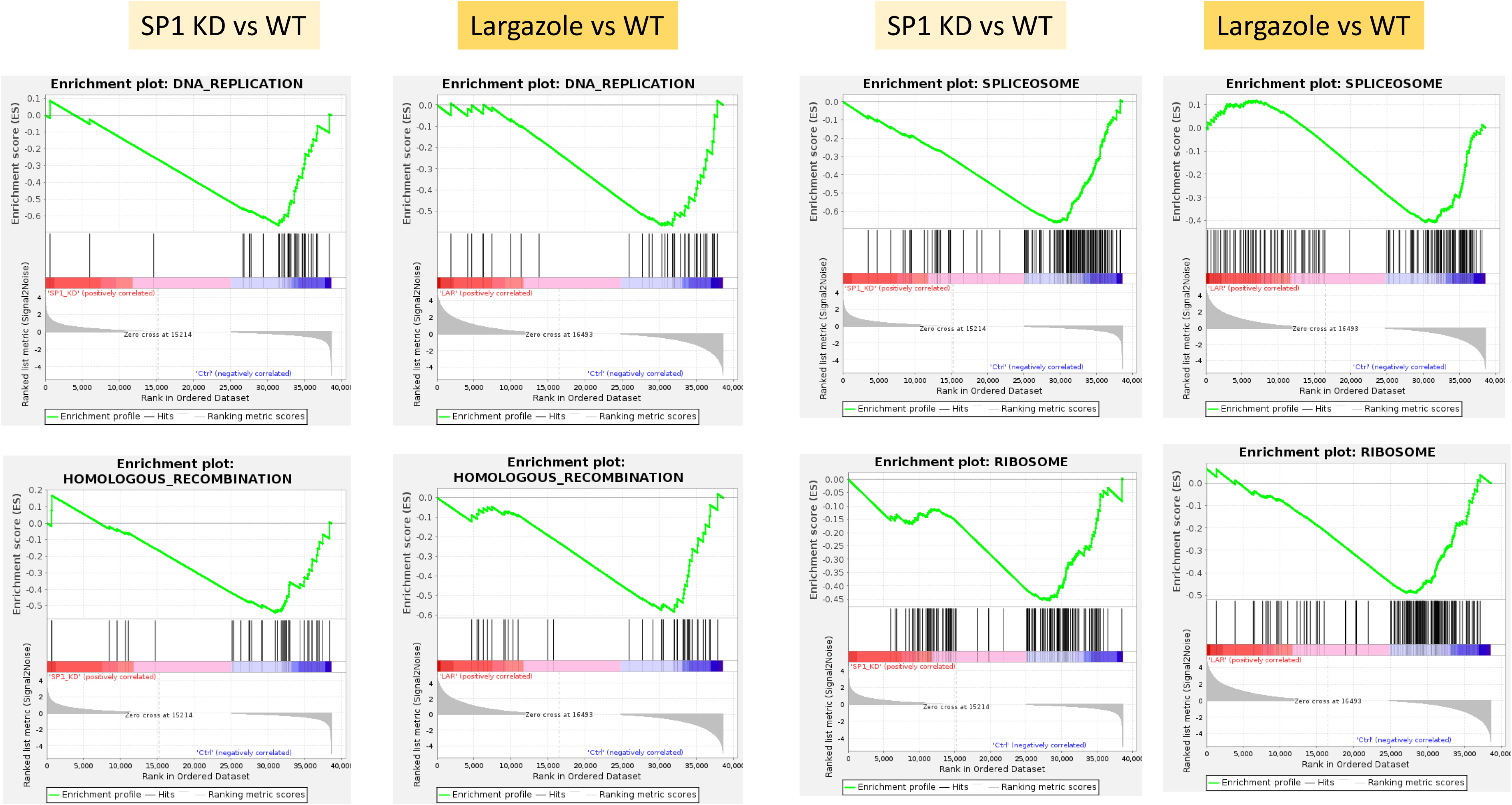
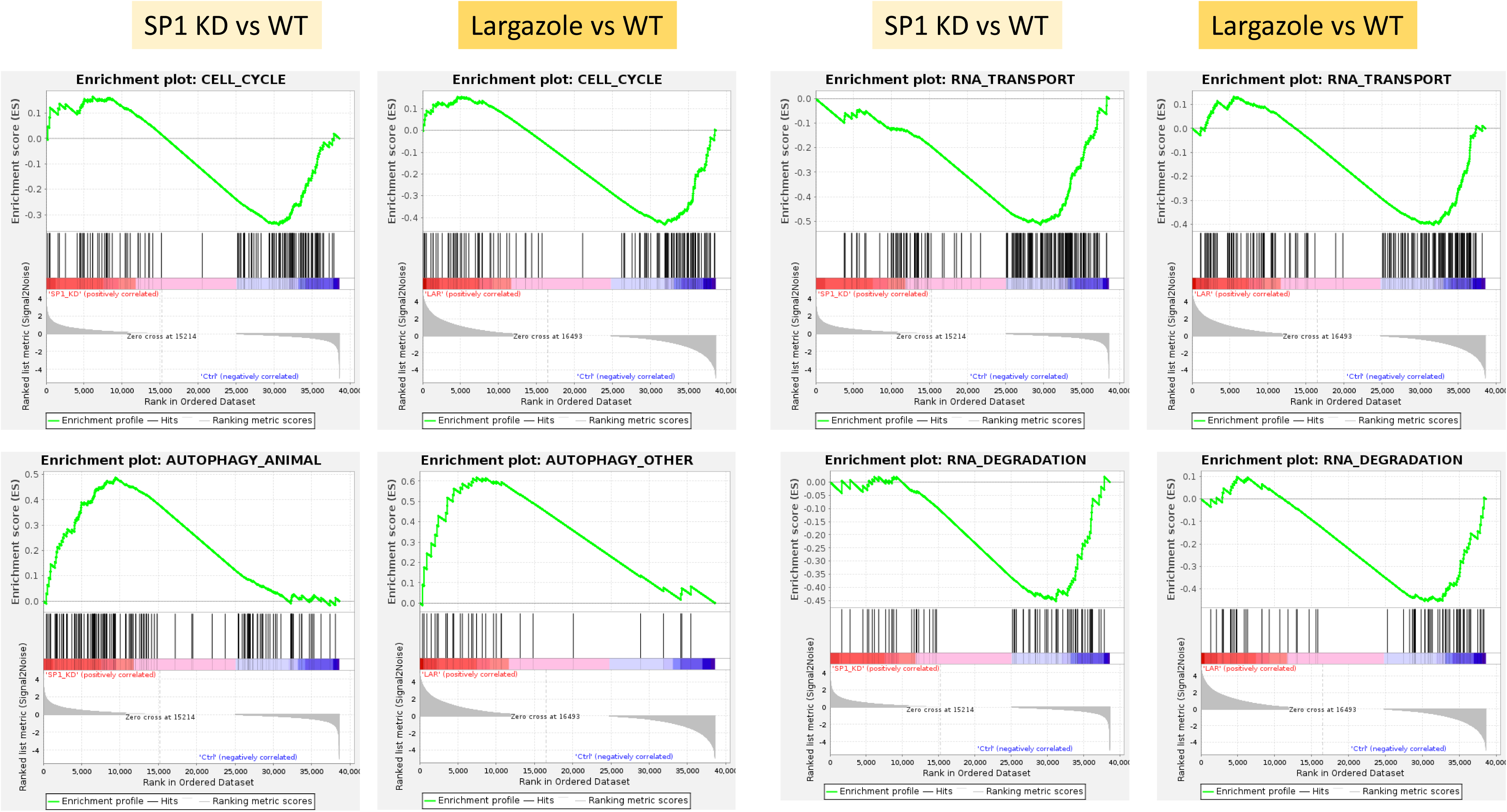
Identification of sets of genes and pathways that are differentially expressed under conditions of SP1 knockdown or largazole treatment by gene set enrichment analysis (GSEA). The size of gene sets and enrichment scores are listed in the tables and enrichment plots for each pathway are shown.

